# Systematic Discovery of Antibacterial and Antifungal Bacterial Toxins

**DOI:** 10.1101/2021.10.19.465003

**Authors:** Nimrod Nachmias, Noam Dotan, Rina Fraenkel, Marina Campos Rocha, Monika Kluzek, Maor Shalom, Arbel Rivitz, Naama Shamash-Halevy, Inbar Cahana, Noam Deouell, Jacob Klein, Neta Schlezinger, Netanel Tzarum, Yaara Oppenheimer-Shaanan, Asaf Levy

**Affiliations:** The Department of Plant Pathology and Microbiology, The Institute of Environmental Science, The Robert H. Smith Faculty of Agriculture, Food, and Environment, The Hebrew University of Jerusalem, Rehovot, Israel; The Department of Biological Chemistry, Alexander Silberman Institute of Life Sciences, Faculty of Mathematics & Science, The Hebrew University of Jerusalem, Jerusalem, Israel; Koret School of Veterinary Medicine, The Robert H. Smith Faculty of Agricultural, Food & Environment, The Hebrew University of Jerusalem, Rehovot, Israel; Department of Materials and Interfaces, Weizmann Institute of Science, Rehovot, 76100, Israel

**Keywords:** Bacterial toxins, polymorphic toxins, inter-microbial competition, bacterial genomics, Antimicrobials, Antifungals, DNAse

## Abstract

Microbes employ toxins to kill competing microbes or eukaryotic host cells. Polymorphic toxins are proteins that encode C-terminal toxin domains. Here, we developed a computational approach to discover novel toxin domains of polymorphic toxins within 105,438 microbial genomes. We validated nine short novel toxins (“PTs”) that cause bacterial or yeast cell death. The novel PTs are encoded by ∼2.2% of the sequenced bacteria, including numerous pathogens. We also identified five cognate immunity genes (“PIMs”) that neutralize the toxins. Intriguingly, we observed an antifungal effect of the PTs against various pathogenic fungi. The toxins likely act as enzymes that cause severe damage to cell shape, membrane, and DNA. Finally, we solved the 3D structure of two PTs in complex with their PIMs, and showed that they function as novel DNAses. The new potent toxins likely play key roles in inter-microbial competition and can be utilized in various clinical and biotechnological applications.

## Introduction

Microbes have evolved a plethora of mechanisms to attack surrounding eukaryotic and prokaryotic cells. One common killing or growth inhibition strategy is the injection of protein toxins through various secretion systems, such as the type III-VII bacterial secretion systems (T3SS-T7SS). Killing competing microbes or host cells can lead to a shift in the microbial community structure or a disease to the host, respectively. The proteins that are being injected target essential cellular components such as the DNA ^1–5^, RNA ^6, 7^, cell membrane ^8^, cell wall ^9, 10^,the translation machinery ^6, 7, 11, 12^, and ATP ^13^. Among the antagonistic molecular tools employed by bacteria, polymorphic toxins ^14, 15^ stand out as an intriguing toxin protein class. These are fairly large proteins, which can surpass 3,000 amino acids (aa) ^15^, that are defined by a specific protein domain architecture (Figure S1). The protein starts with an N-terminal domain that serves as a trafficking domain, which associates with a secretion system or serves as a signal peptide translocating the protein to the periplasm. The trafficking domain is followed by repetitive elements that might stretch over significant length, such as Rearrangement Hot-Spot (Rhs) or hemagglutinin repeats. Toward the C-terminal end of the protein there are releasing peptidase or pre-toxin domains. The C-terminus of the protein is a toxin domain that is eventually delivered into and kills target cells by forming pores in the cell envelope, degrading or blocking the synthesis of phospholipids, peptidoglycan, and nucleotides, or degrading nucleic acid derivatives ^14^. The toxin encoding genes are usually followed by immunity genes that protect the toxin producing microbe from self-intoxication (Figure S1) ^16^. The name ‘polymorphic toxin’ is derived from the modular nature of these systems, which display high polymorphism in all of its constituent domains, including at the toxin position. Namely, polymorphic toxin variants can exist in different organisms, with some domain architectural changes which allow these proteins to be delivered by different secretion systems, to be processed differently, and to kill cells using a variety of mechanisms.

Polymorphic toxins mostly play important antagonistic roles in various microbes. For example, Tc toxins are known for their insecticidal role. A recent survey of TcC proteins of bacterial Tc toxins identified that these are polymorphic toxins with over 100 different putative toxic domains ^17^. Firmicutes in the human gut are armed with LXG polymorphic toxins which mediate antimicrobial antagonism ^10^. *Staphylococcus aureus* employs T7SS to secrete a polymorphic toxin with a nuclease activity to kill rival bacteria ^18^ and Bacteroidetes likely use T9SS to secrete antibacterial DNase polymorphic toxin ^19^. *Burkholderia thailandensis* uses contact-dependent growth inhibition (CDI) and associated delivered polymorphic toxins to antagonize foreign bacteria and to promote biofilm formation in resistant kin cells ^20^. The soil microbe *Serratia marcescens* secretes Rhs-encoding effectors with DNAse activity to mediate intraspecies competition ^4^. *Aeromonas dhakensis* secretes, via T6SS, a polymorphic toxin that acts as an endonuclease ^5^. Myxobacteria use arrays of polymorphic toxins that are transferred through an outer membrane exchange (OME) mechanism to discriminate clonal cooperators versus non-self and to ensure populations are genetically homogenous ^21, 22^.

Given their highly conserved functions in inter-microbial antagonism, it is clear that discovery of novel polymorphic toxins will provide a better mechanistic understanding of microbial colonization in various habitats, and it may lead to elucidation of novel efficient antimicrobials, as these toxins evolved to kill target cells in low concentrations. Thus, systematic discovery of novel toxins, and characterization of their modes of action and targets may promote development of novel therapies for medical use ^23^ and biotechnological applications such as base editing ^24^. Notably, the genomic revolution has uncovered vast genetic space from understudied non-model organisms, which also presumably cope with bacterial and fungal competitors and therefore evolved yet unknown polymorphic toxins. Bioinformatic mining of these genomes followed by experimentation can lead to the discovery of toxins and new modes of action.

Here, we performed a large-scale discovery followed by an in depth experimental characterization of new toxin domains that are abundant in microbial polymorphic toxins. First, we developed a computational framework to predict novel C-terminal toxin protein domains based on their association with polymorphic toxins across 105,438 microbial genomes. The association of these putative toxin domains with different trafficking domains suggests that they are employed as weapons secreted by various bacterial secretion systems. For several toxin domains, we also identified adjacent putative cognate immunity genes that are syntenic with the toxin domain. We focused on novel toxin domains that lacked amino acid sequence similarity to known toxins. Then, heterologous expression experiments of toxin candidates were performed to test their cellular toxicity. Our experiments uncovered nine new toxin domains that efficiently kill *E. coli* or *S. cerevisiae*. We found that the novel toxins are encoded by ∼2.2% of sequenced bacteria, including major human, animal, and plant pathogens. For five toxins cognate immunity proteins were identified and rescued the expressing cells from toxicity. Intriguingly, the toxins demonstrated potent antifungal and antibacterial properties when mixed with pathogenic fungi or with Gram-positive bacteria, and they could penetrate fungal cells. We further used fluorescence microscopy and observed various cell morphology damage types that are caused by the activity of toxins with diverse modes of action. Furthermore, we revealed and mutated amino acids that are critical for cellular toxicity, and are predicted to be located within the toxin catalytic sites. Finally, we solved the 3D structure of two novel toxin-immunity protein complexes and experimentally demonstrated that the toxins have efficient DNAse activity. Hence, this work significantly expands our knowledge of conserved and potent microbial toxins that are used in warfare against prokaryotes and eukaryotes, and it elucidates mechanistic details regarding activity of the novel toxins. Importantly, these toxins can serve as the basis for development of new antibacterial and antifungal treatments.

## Results

### *In silico* discovery of novel C-terminal toxin domains (PTs) of polymorphic toxin proteins and their cognate immunity genes (PIMs)

Our objective was to discover novel toxin domains that are located at the C-terminal end of polymorphic toxin proteins. The algorithm we developed is described in Figure 1A. First, we compiled a list of 217 potential polymorphic toxin domains, defined as trafficking, repeat, pre-toxin or toxin domains, by performing an exhaustive iterative search in the Pfam database (Table S1, Methods). Using these domains, we searched for polymorphic toxin genes that encode these domains within 105,438 microbial genomes. From these polymorphic toxin genes, we retrieved their putative C-terminal toxic domains (serving as toxin candidates), defined as the last 133 aa of the protein, which is the median size of a protein domain in Pfam ^25^. Next, we clustered these 2.6 million sequences to yield 152,150 clusters representing putative toxin families. To reduce false positive hits, several filters were applied to the clusters (Figure 1A). We kept clusters that had no hits to known domains, were sufficiently taxonomically diverse, and were found in high diversity of polymorphic toxin protein architectures (Methods). Finally, we obtained a list of putative C-terminal domains of PTs that very likely serve as potent, novel toxins (Tables S2-S3).

**Figure 1.**
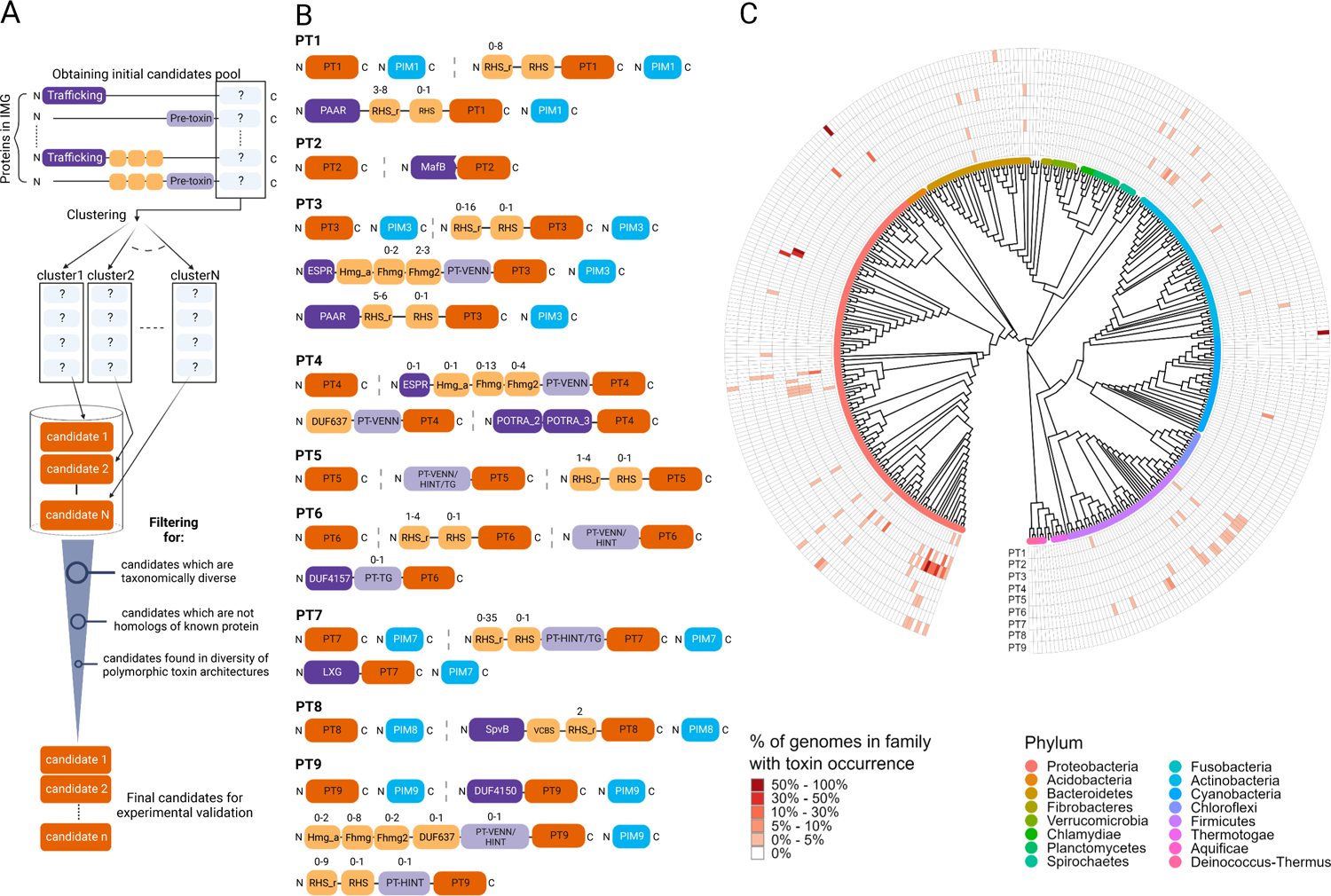
Computational prediction of novel toxin domains (PTs) and cognate immunity genes (PIMs) in polymorphic toxin proteins. **(A)** A pipeline for prediction of novel toxic protein domains (PTs). C-terminal sequences of predicted polymorphic toxin proteins having one or more trafficking/pre-toxin/repeat domains were considered as potential toxin domains (PTs). These C-termini were clustered based on amico acid sequence similarity to obtain an initial pool of toxin candidates. Candidates were filtered out in multiple stages, keeping ones which are sufficiently taxonomically diverse, novel (not found by a sensitive sequence similarity-based search), and found in high diversity of polymorphic toxin protein architectures. This pipeline led to a final list of domain candidates that very likely bear toxic activity. For each final candidate, we obtained a representative by searching for a single domain gene which is a homolog of the cluster members. The representatives were used in the experimental validation step. **(B) Most abundant protein architectures that include our novel PTs.** The protein domains were annotated with Pfam domains using hmmscan (RHS_r, Hmg_a, Fhmg, and Fhmg2 stand for RHS_repeat, Haemagg_act, Fil_haemagg, and Fil_haemagg_2, respectively). The toxin domains (PTs) are shown in orange, cognate adjacent immunity genes are in blue, repeat domains are in beige, pre-toxin domains are in light purple, and trafficking domains are in dark purple. The number above the domain is the range of possible consecutive occurrences of the domain in this type of domain architecture. Exact numbers of occurrences of the architectures are shown in Table S4. **(C) Taxonomic distribution of the novel toxins.** The phylogenetic tree was constructed using marker genes from representative genomes from each taxonomic family. Therefore, each leaf in the phylogenetic tree represents a bacterial family. Leaf color represents the phylum of the family and outer rings represent the percentage of genomes in the matching family that contain at least one occurrence of the toxin, according to the IMG database (shades of red). A phylogenetic tree with family names is presented in Figure S2.

We experimentally validated nine novel toxin domains, termed “PT1-9” (an acronym for <u>P</u>olymorphic <u>T</u>oxin C-terminal domains) (Figure 1B, Tables S2, S4-S5). Notably, we observed that the novel PTs are part of varied large polymorphic toxin proteins that have a large diversity of N-terminal trafficking domains (Figure 1B). The trafficking domains can lead to PT secretion through various secretion systems. Therefore, we predict that PT1, PT3, and PT9 are secreted through the T6SS as they are fused to PAAR ^26^ and DUF4150 ^27^ domains, PT6 is secreted through the extracellular contractile injection system (eCIS) as it is fused to DUF4157 eCIS marker domain ^28^, and PT4 is secreted via the T5SS as it is fused to Extended Signal Peptide of Type V secretion system (ESPR) domain ^29^. In addition, PT7 is likely secreted via the T7SS as it is fused to LXG domain ^10^, and PT4 is likely secreted through the Omp85/TPS β-barrel proteins as it fused to POTRA_2 and POTRA_3 domains ^30^. Importantly, for certain PTs we identified a cognate downstream gene that we predicted to serve as an immunity gene (Figure 1B, Tables S2). We termed these genes <u>P</u>olymorphic <u>Im</u>munity genes (“PIMs”). The PT-PIM pairs were predicted based on conserved synteny of the two gene families across numerous genomes (Tables S4-S5).

### Distribution of the novel toxins across bacterial taxa, ecosystems, and in bacterial pathogens

To provide a better understanding of the broad function of the novel PTs we analyzed their taxonomic distribution (Figures 1C, S2) and their presence in various bacterial pathogens (Table S6). PT1 is found in Proteobacteria, Firmicutes and Bacteroidetes phyla including in the human pathogens *Clostridium botulinum*, *Burkholderia cenocepacia*, *Cronobacter sakazakii*, and the plant pathogen *Dickeya zeae*. PT2 is Neisseriaceae-specific, encoded in the human pathogens *Neisseria gonorrhoeae* and *Neisseria meningitidis*. PT3 is widely distributed along five different phyla: Firmicutes, Actinobacteria, Fusobacteria, Bacteroidetes, Proteobacteria. It is found in the human pathogens *Pseudomonas aeruginosa* (Table S7), *Klebsiella aerogenes*, *Salmonella enterica*, *Acinetobacter baumannii*, and *Cronobacter sakazakii* and the plant pathogen *Ralstonia solanacearum*. The PT3 representative we validated from *Ralstonia solanacearum* (see below) shares weak amino acid sequence similarity (41% identity) with the C-terminal of the T6SS effector RhsP2 from *Pseudomonas aeruginosa* ^31, 32^ and likely these toxins belong to the same toxin superfamily. PT4 is found only in the Proteobacteria phylum including in the human pathogens *Klebsiella aerogenes*, *K. pneumoniae*, *K. variicola*, *Yersinia pestis*, *Y. pseudotuberculosis*, *Serratia marcescens*, *Salmonella enterica*, *Cronobacter sakazakii* and in the animal and plant pathogens *Edwardsiella anguillarum*, *E. ictaluri*, *Photorhabdus luminescens*, and *Pantoea agglomerans*. PT5 is found in Actinobacteria, Cyanobacteria and Proteobacteria phyla, including in the human pathogen *Vibrio vulnificus*. PT6 is found in Firmicutes, Actinobacteria, Spirochaetes, Bacteroidetes and Proteobacteria phyla but is absent from pathogens. PT7 is found in Firmicutes and Proteobacteria phyla, including the human pathogens *Bacillus cereus*, *Acinetobacter baumannii*, *Listeria monocytogenes* and in the insect pathogen *Bacillus thuringiensis*. PT8 is specific to the Leptospiraceae family, found almost exclusively in the causing agent of Leptospirosis, *Leptospira interrogans*. PT9 is found in Cyanobacteria, Bacteroidetes and Proteobacteria phyla. It is found in the human pathogen *Bordetella genomosp* and the insect pathogen *Xenorhabdus cabanillasii*. We noted that Proteobacteria phylum encoded all nine novel PTs except for PT8, with Enterobacteriaceae, Morganellaceae, Pseudomonadaceae, and Neisseriaceae families encoding four PTs each. Interestingly, although Cyanobacteria is a relatively sequenced phylum with thousands of available genomes we identified a single new PT family (PT5) in a single cyanobacterial family, Microcoleaceae.

We also correlated the toxin encoding bacteria with the bacterial habitat, life-style, and physiology suggesting that toxin presence might be associated with these biological features (Table S7). This can allow us to infer important ecological features that likely pertain to the toxin activity. We statistically corrected this analysis for phylogenetic biases in the genomic data, as the PTs can be found in phylogenetically related genomes which are over-represented in sequence databases (Brynildsrud et al. 2016) (Methods). Three PTs, PT3, PT4, and PT9, are enriched in host-associated bacteria. PT4 is enriched in motile bacteria isolated from humans, fish, insects, and nematodes. PT3 is enriched in specific human-associated niches where bacteria dwell including sputum, the excretory system and the respiratory system. Interestingly, PT5 is enriched in environmental bacteria, such as the aquatic *Rheinheimera baltica* and *Cupriavidus pauculus,* and the plant-associated *Cupriavidus plantarum*. Overall, the nine novel PTs are widespread across the microbial world, and are encoded in microbes isolated from various habitats, of different life-styles, including in major human, animal, and plant pathogens.

### Experimental validation of novel PTs shows that they efficiently kill bacteria or yeast

To experimentally test if the predicted toxic domains (PTs) will exert a toxic phenotype in bacterial or eukaryotic cells, we performed heterologous expression assays in *Escherichia coli* and in *Saccharomyces cerevisiae*. Since we could not define the precise PT domain borders within their encoding proteins (Fig 1B), we searched for short, single domain genes that were homologous to the PTs, and selected them as representatives for each PT family (Tables S2 and S3). The selected short genes were taken from various Gram-positive and -negative genomes, and the genes encoded putative toxin domains shorter than 150 aa that were not mapped to any known Pfam domain. Our selection of *E. coli* or *S. cerevisiae* as the PT target cells was partially based on presence or absence of predicted cognate immunity genes next to the PTs (prokaryotic-targeting toxins likely have presence of immunity genes). We used codon-optimized, *de novo* gene synthesis to clone the putative antibacterial PTs into pBAD24 plasmid and expressed them in *E. coli* with arabinose induction and the anti-eukaryotic PTs in pESC plasmid in *S. cerevisiae* with galactose induction. As controls we used empty vectors. Strikingly, already in the synthesis and cloning stage we noticed that three genes, despite being synthesizable, were toxic and recalcitrant to cloning and propagation in plasmids suggesting that even leaky expression was sufficient for killing the hosting microbe. Genes that were successfully cloned in *E. coli* and *S. cerevisiae* vectors were plated on agar plates supplemented with inducer (arabinose or galactose) or a repressor as control (glucose). Importantly, nine genes (PT1-9) exhibited high toxicity to *E. coli* of up to four orders of magnitude reduction in CFU (Figures 2A and S3A). PT4 and PT6 were not toxic in the cytoplasm so we introduced an N terminal periplasmic translocation signal (twin-arginine translocation signal) and confirmed their toxicity in the bacterial periplasm(Figures 2B and S3B). Moreover, the PT activity of nearly all toxins inhibited growth in liquid media (Figure 2C). We monitored a rapid growth arrest resulting in a range of 4-10 fold longer lag time or approximately two-fold decrease in exponential growth bacterial concentrations, as measured by optical density, of PT-expressing strains compared to the control. Remarkably, PT3, PT6, and PT7 were toxic to both yeast and bacterial cells (Figures 2A-D and S3C). These findings demonstrate that nine novel toxins are highly toxic to recipient bacterial and eukaryotic cells.

**Figure 2.**
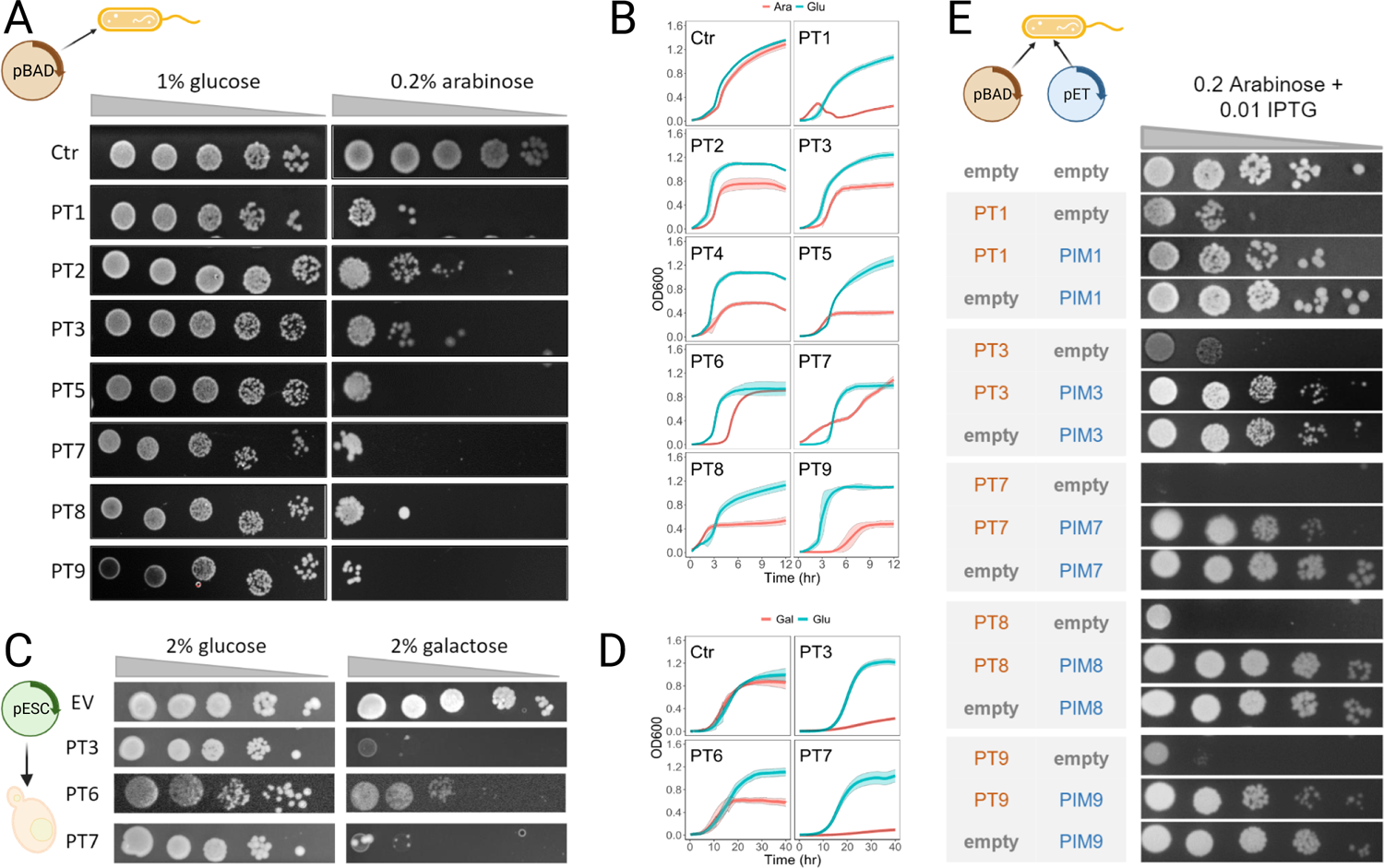
Novel PTs efficiently kill bacteria and/or yeast and PIMs neutralize the toxicity. **(A)** The toxins were cloned into *E. coli* BL21 (DE3) under the control of an arabinose-inducible promoter. As control, cells that lack the toxin (empty vector) were cloned (Ctr). Bacteria were plated in 10-fold serial dilution on LB-agar in conditions that repress gene expression (1% glucose) or induce expression (0.2% arabinose). **(B)** Growth curves of *E. coli* BL21 (DE3) in liquid culture. Cells were grown in LB-liquid media supplemented with 1% glucose or 0.2% arabinose. OD600, optical density at a wavelength of 600 nm. (n=3, ± SD for values). **(C-D)** Several PTs were cloned into *Saccharomyces cerevisiae* BY4742 under the control of a galactose-inducible promoter. As a control, cells that lack the toxin (empty vector) were cloned (Ctr) **(C)** Yeast cells were plated in 10-fold serial dilution on SD-agar in conditions that repress gene expression (1% glucose) or induce expression (1% galactose). **(D)** Growth curves demonstrate the toxic effect of PT3, PT6, and PT7 toxins on *S. cerevisiae* BY4742 growth for 40 hours. Cells were grown in SD-liquid media supplemented with 2% glucose or 2% galactose. OD600 of technical replicates were recorded (n=3, ±SD for values). **(E)** PIMs provide protection against toxicity of cognate PTs. The toxins (PT) were cloned under the control of an arabinose-inducible promoter (pBAD24) and the protein immunity protein (PIM) were cloned under the control of an inducible promoter system that is active only in the presence of IPTG (pET28) into *E. coli* BL21 (DE3). The plates included both inducers.

### Toxicity of PTs is neutralized by cognate immunity genes (PIMs)

Antibacterial toxins encoded by bacteria are often escorted by adjacent immunity genes, as known for T6SS, toxin-antitoxin, and contact-dependent inhibition systems ^16, 33–35^. We identified five cognate PIM genes that are found in synteny with the toxin domain, thus predicted to serve as immunity genes (Figure 1B, Table S2). We tested whether PIM genes can neutralize their cognate PTs by co-transformation and co-induction of the putative PT-PIM pairs carried on different expression vectors, pBAD24 and pET28a for PT and PIM, respectively. As control, *E. coli* cells were co-transformed with empty vectors along with the PT or the PIM. Bacteria harboring the PIM grew similarly to the control strain on solid media and exhibited normal cellular morphology showing that the five PIMs are non-toxic to *E. coli* (Figures 2E and S3D). As predicted, these PIMs successfully rescued the bacteria from intoxication by their cognate PTs on solid agar and in liquid media (Figure 2E). Moreover, using fluorescence microscopy, we observed that expression of PTs led to cellular aberrations at the single cell level, while expression of PT-PIM pairs led to morphologically normal cells, again demonstrating that PIM expression inhibits the effects of PT expression (Figure S4). These cellular aberrations will be discussed in further detail in a later section. We note that the novel PIMs lack sequence similarity to any protein of known function (Table S2).

### Conserved amino acids and protein structure prediction suggest involvement of the toxins in enzymatic activities

A well-established strategy in bacterial competition as well as in virulence in eukaryotes is the use of highly active enzymes as cytotoxic effectors ^15, 36–38^. Thus, we sought to explore the mechanism of action of the newly discovered toxins. First, we characterized the folded toxin structures and their conserved residues that are required to execute their lethal functions. To explore the folding of the proteins we predicted the PT 3D structures using roseTTAFold ^39^ and compared the structures to known structures using DALI ^40^. Our results suggest that most of the PTs we discovered represent novel protein structures (Table S9). However, PT3 predicted structure resembles the Diphtheria and Cholix toxins which act as ADP-ribosyltransferase of EF-2 protein ^41, 42^, and PT4 resembles proteins that act as NADH dehydrogenase suggesting that the PTs maintain at least somewhat similar enzymatic activity. The other PTs presented weaker structural similarities to known proteins (lower Dali Z-scores) or inconsistent hits. Next, we searched using two approaches for conserved residues that are critical for toxin activity and likely serve in potential enzymatic activity. First, for each PT we generated HMM logos based on alignments of all the toxin homologs and selected the most conserved residues (Figures 3A and S4). Second, we applied Jensen-Shannon Divergence scoring (Methods) to each position to infer functionally important residues from sequence conservation ^43^. Among the high scoring positions, we looked for residues with high propensity to serve in catalytic sites of enzymes, i.e. residues with basic, acidic or nucleophilic nature ^44^. Using these features, we selected 1-3 residues to mutate per toxin. We substituted the conserved residue with alanine via directed mutagenesis of the pBAD24 plasmid harboring the toxins. Strikingly, point mutations of selected PTs led to complete abolishment of the toxic phenotype, restoring normal growth when bacterial cultures were grown on solid media (Figure 3B) and on liquid media (Figure S6). We observed normal or nearly normal cellular morphology at the single cell level of the mutants in comparison to the wild-type toxins (Figure S6).

**Figure 3.**
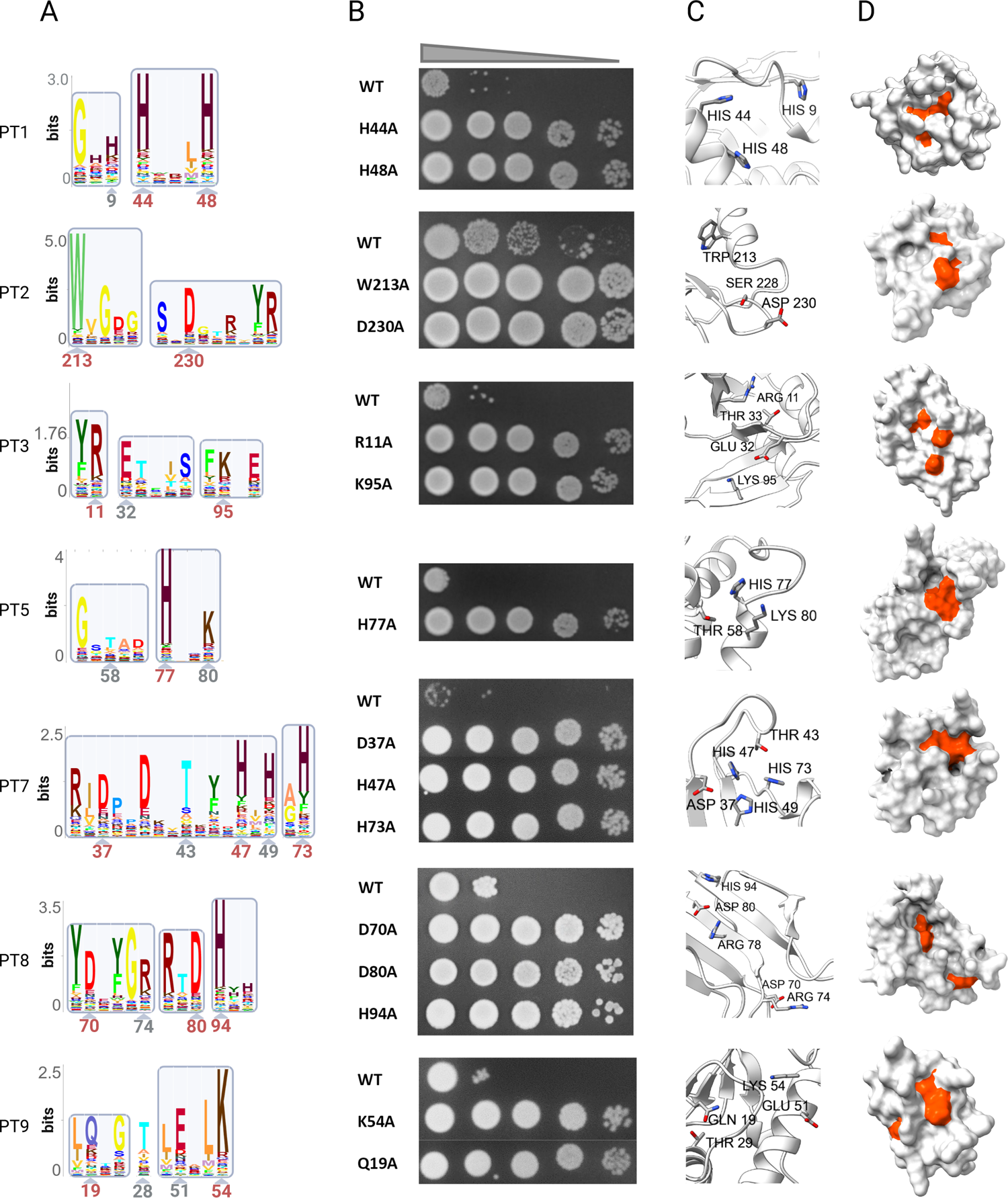
Critical residues and predicted protein structures of the novel PTs. **(A)** HMM logo of conserved boxes in the novel toxins (names appear from top to bottom on the left side of the panel). Highly conserved regions in logos generated by skylign were cropped and are numbered by position with respect to each experimental candidate. Position numbers marked in red note experimentally tested residues. **(B)** The toxins (WT) and generated toxin mutants were cloned into *E. coli* BL21 (DE3) under the control of an arabinose-inducible promoter. Bacteria were plated in 10-fold serial dilution on LB-agar in conditions that induce expression (0.2% arabinose). **(C)** Microenvironment 3D modeling of roseTTA Fold predictions for candidates (from top to bottom; % confidence): PT1 (72%), PT2 (52%), PT3 (84%), PT5 (79%), PT7 (85%), PT8 (50%) and PT9 (66%). Backbone is displayed as a cartoon model. Residues with high conservation ratios and propensity for involvement in catalysis are displayed as sticks without hydrogen, residue three letter code and position number are marked. **(D)** Surface displays of the predicted models. Residues that were marked in **(B)** are colored in orange red.

**Figure 4.**
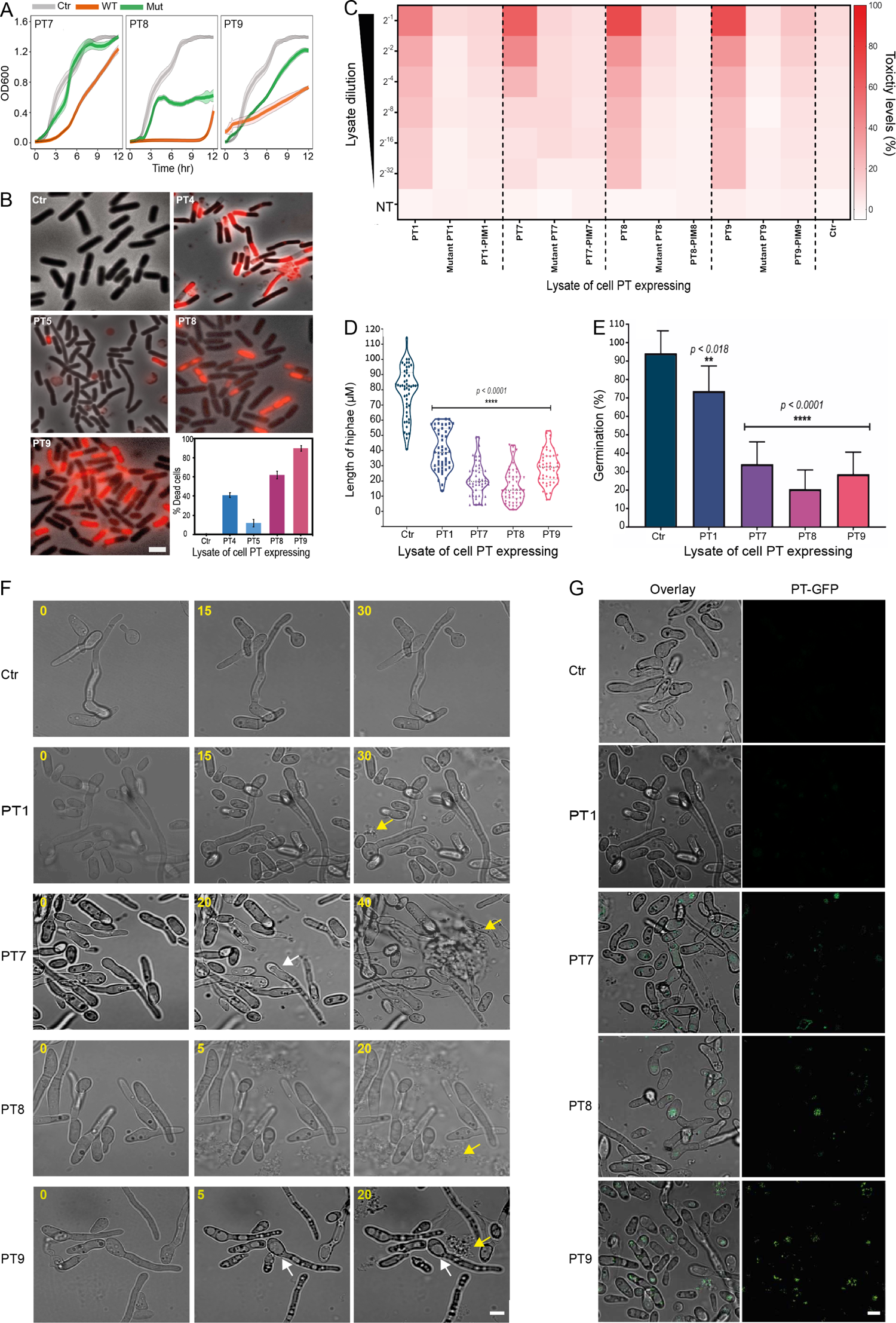
Antibacterial and Antifungal activities of PT toxins. (**A**) and (**B**) *B. Subtills* (NCIB 3610) were grown in LB-liquid media supplemented with lysates of PT expressing *E. coli* cells. Ctr, control of lysates of *E. coli* cell PT expressing with an empty vector. **(A)** Growth curves of *B. Subtills* (NCIB 3610). OD600, optical density at a wavelength of 600 nm. (*n*=3, ± SD for values). **(B)** *B. Subtills* cells were incubated for 2 hours with cell lysates, *s*tained with PI and visualized by fluorescence microscopy. Shown are overlay images of phase contrast (gray) and fluorescence from PI staining (red). Scale bar corresponds to 2 µm. **(C)** *F. oxysporum* Fol4287 was grown with lysates of PT-expressing *E. coli* cells. The heat map represents toxicity levels that are computed based on growth inhibition. To determine the effect of PTs on fungal growth the indicator Alamar blue Fluorescence intensities (FI) of treated and untreated samples for each strain were obtained after 48 hours of incubation. FI were used to calculate the percentage of toxicity. PT1, PT7, PT8, PT9 - fungal toxicity when mixed with lysates of *E. coli* cells expressing wild-type PTs, Controls: PT mutants and PT-PIM gene pairs. Ctr; lysate of *E. coli* cells with an empty vector. NT: Non treatment. **(D)** Measurements of *F. oxysporum* Fol4287 hyphal growth following treatment with different PT-expressing *E. coli* lysates. (**E**) Germination level of *F. oxysporum* Fol4287 following treatment with different PT-expressing *E. coli* lysates. (**F**) Damage caused to fungal cells. Images from time-lapse microscopy. *F. oxysporum* Fo1428 cells incubated with different PT-expressing *E. coli* lysates. Capture time is indicated in the top left corner. Arrows point to vacuole modifications (white) and degraded cells (yellow). (**G**) PTs penetrate fungal cells. Overlay of brightfield and green (left) and PT-GFP (right) expression images of the *F. oxysporum* Fol4287 fungal cells treated with lysates of E. coli cell expressing PT1 and PT7 (after 30 min of incubation) or PT8 and PT9 (after 5 min of incubation). Scale bars in F and G correspond to 5 µm. In all figures Ctr is lysate of *E. coli* cells with an empty vector.

Next, we mapped the residues that were critical to PT function to the folded protein. Surprisingly, highly conserved residues with high propensity to participate in catalysis were spatially clustered and directed towards each other on the simulated atomic microenvironment (Figures 3C and S7). Surface display of the model shows that these residues are generally found in proximity of pocket-like intrusions on the protein surface (Figure 3D) which is a typical trait of enzymes. Taken together, these data suggest that the new PTs serve as a diverse group of enzymatic toxins.

### Lysates of PT expressing cells demonstrate potent antibacterial and antifungal activity against *Bacillus subtilis* and various pathogenic fungi

We next searched for potential extracellular antimicrobial activity against bacteria and fungi. Unfortunately, we could not purify the wild-type PTs due to their high toxicity to the producing cells. As a substitute, we used cell lysates of *E. coli* expressing PTs and empty vector controls against a set of target organisms including *Bacillus subtilis*, *E. coli*, and six pathogenic fungi species that cause diseases to humans, animals or plants (Table S11). The target cells were grown in media supplemented with the bacterial lysates of all PTs. We observed that *B. subtilis* growth in liquid media was fully inhibited by PT8, and was partially inhibited by PT7 and PT9 (Figure 4A). Lysates of the PT mutants led to reduced toxicity and lysates of *E. coli* expressing an empty plasmid (Ctr) were non toxic (Figure 4A). Furthermore, we quantified cell death based on cellular penetration of propidium iodide stain, which enters cells that have a damaged membrane^45^. *B. subtilis* cells that were incubated for two hours with cell lysates of PT4, PT5, PT8 and PT9 expressing cells showed sharp increase in dead cells (Figure 4B). We stained the *B. subtilis* membrane and DNA and subjected to analysis by fluorescence microscopy. While *B. subtilis* cells treated with Ctr lysates had normal morphology, cells treated with PT-expressing cell lysates demonstrated cell division inhibition (PT1 and PT4), disintegration of the membrane (PT4), abnormal membrane and abnormal cell sizes (PT8), and partial cell rounding (PT9) (Figure S8). When *E. coli* cells were incubated with PT-expressing cell lysates, no killing was observed (not shown).

Remarkably, lysates of different PT-expressing cells were toxic at different levels to the six species of pathogenic fungi we tested, including species of *Aspergillus nidulans*, *Aspergillus fumigatus, Candida albicans*, *Candida auris, Cryptococcus neoformans*, and *Fusarium oxysporum*. The activity was concentration-dependent, as toxicity decreased when lysates were serially diluted (Figures 4C and S9). Moreover, the toxicity was abolished or decreased when the fungal cells were grown with lysates of cells expressing PT mutant, PT-PIM complex or an empty plasmid (Figures 4C and S9). Surprisingly, each of the toxins we tested demonstrated antifungal activity against at least one fungal strain (Figure S9). PT1, PT2, and PT9 were toxic against nearly all the strains at their highest concentrations (Figures S9A-B and F). PT3 was the least toxic, yet still had ∼50% toxicity against *C. auris* and *C. neoformans* at its highest concentration (Figure S9C). PT7 showed ∼80% toxicity against *A. nidulans* and *C. albicans* at its highest concentration (Figure S9D), and PT8 demonstrated 100% toxicity against *A. fumigatus* and maintained >60% toxicity even in 2^-4^ dilution (Figure S9E). To further analyze the antagonistic effect on fungi, we focused on the PTs antifungal effect on *F. oxysporum*, the causal agent of vascular wilt disease in plants and an emerging opportunistic human pathogen ^46, 47^. Lysates of cells expressing PT1, PT7, PT8, and PT9 reduced hyphal length by at least 50% (Figure 4D) and PT7, PT8, and PT9 reduced germination by over 67% in comparison to Ctr (Figure 4E). We examined the consequence of growing *Fusarium* with the PT expressing cell lysates using time-lapse fluorescence microscopy. Remarkably, we observed that a short time after supplementing the lysates, the fungi start degrading, e.g. as short as five minutes after adding PT8 (Figure 4F). As part of degradation, we could observe formation of cytoplasmic vacuoles when the fungi were incubated with lysates of PT8- and PT9-expressing cells (Figure 4F). To determine if the toxins could enter *F. oxysporum* cells, we fused green fluorescent protein (GFP) to the C terminus of PTs and incubated the lysates of PT-GFP expressing cells with the fungi. We followed the toxins under the microscope and found that the toxins clearly penetrated the cells in up to 30 minutes (Figure 4G). Taken together, these results demonstrate that each toxin tested has the ability to kill different bacteria or fungi when supplemented extracellularly.

### The effect of the novel PTs activities on cell size, cell membrane and DNA morphology

Following the elucidation of new PTs that kill cells intracellularly and extracellularly, we studied more closely the mode of action of the PTs. We examined the phenotypic effect caused by the novel PTs to the transformed cells at a single cell resolution. PT-expressing cells were stained with membrane and DNA stains and visualized by fluorescence microscopy at approximately 40 minutes post-induction (Figure 5). The results suggest that some of the toxins cause direct or indirect damage to the DNA. Expression of PT1, PT3, and PT8 led to a ‘cloud’ of degraded DNA outside the cells. PT3 and PT7-expressing cells retained their characteristic wild-type rod shape and membrane architecture, but their chromosomes appeared to be contracted, and in some cells had vanished completely (the signal from DAPI staining was hardly detectable), suggesting that PT3 and PT7 may degrade the chromosome. Remarkably, PT9 expression led to mini-cells and chromosomal condensation and DNA foci at the cell center which is reminiscent of nalidixic acid treated cells ^48^. Other toxins likely target the cell membrane or the cell wall. For example, when we expressed PT4 in the periplasm the membrane disintegrated and the cells were swollen and lost their rod-like shape. Induction of PT5 led to several round cells suggesting that this toxin damages the peptidoglycan layer and thereby leading to loss of the rod-like structure. In addition, we observed large cell aggregates. Expression of two other PTs led to cell division inhibition. PT2 induction led to cell elongation and long interconnected cells. Expression of periplasmic PT6 led to massive cell filamentation with multiple nuclei which indicates a strong cell division inhibition. PT8 expressing cells had odd shapes with occasional membrane accumulation. In line with those observations, the different morphological abnormalities may provide directions to more thoroughly study modes of PT-induced cellular toxicity.

**Figure 5.**
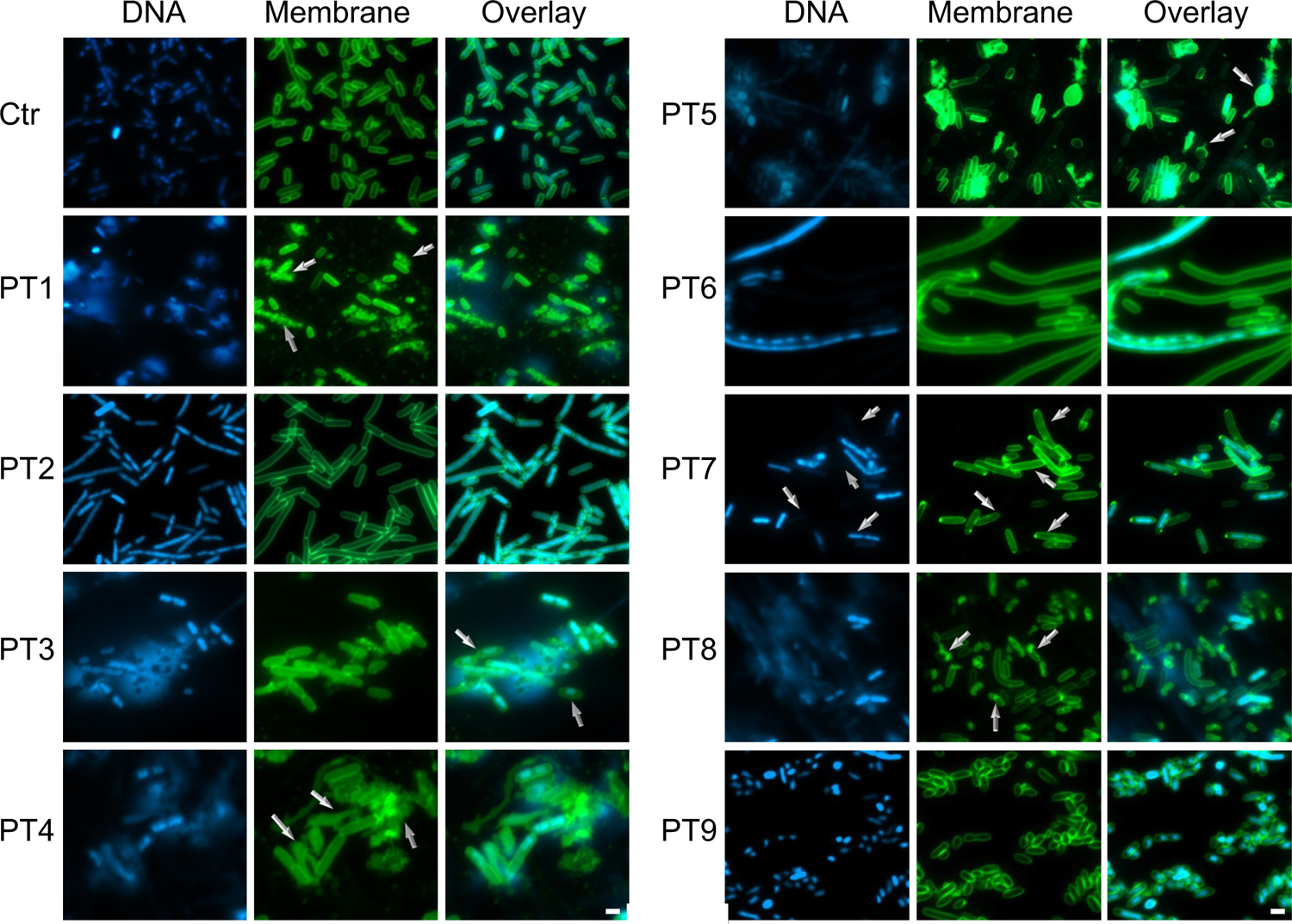
PTs lead to cell death in various manners. *E. coli* BL21 (DE3) cells harboring toxin genes (PT) or empty vector as control (Ctr) were grown in LB media in conditions that induce expression (0.2% arabinose). After 40 min of incubation, samples were stained with membrane and DNA stains and visualized by fluorescence microscopy. Shown are DNA (DAPI) (blue), membrane (FM1-43; green) and overlay images of the bacterial cells with toxins as indicated. Arrows point to abnormal cells; PT3: compressed chromosomes, PT4: membrane disintegration and swollen cells, PT7: missing DNA and membrane movement to the poles, PT8: odd shaped cells. Representative images from a single replicate out of three independent replicates are shown. Scale bar corresponds to 2 µm.

To determine whether some PTs directly affect bacterial membranes, we tested *in vitro* the effect of different PT-expressing cell lysates on large unilamellar vesicles (LUVs) composed of *E. coli* lipid extract with encapsulated calcein. We observed that PT4, PT8 and PT9-expressing cell lysates induced LUVs permeabilization and caused a rapid release of the internal calcein (Figure S10). It is yet unclear what is the exact mode of action of these toxins.

### Biochemical and structural characterization of PT1 and PT7

To conduct biochemical and structural studies, a considerable amount of highly purified protein is required. Initial cytosolic and periplasmic expression experiments of His-tagged wild-type PT1 and PT7 in bacterial cells resulted in undetectable expression of the PTs. Thus, we tested the expression of PT1 and PT7 point mutants (Figure 3) for structural studies. Moreover, PT1 and PT7 point mutants can also be used for functional characterization of the PT1 and PT7 modes of action if the mutants do not fully abolish the enzymatic activity. In parallel, we expressed the immunity proteins PIM1 and PIM7 and co-purified the PT-PIM complexes.

Based on the phenotypic characterization (Figure 5), we hypothesized that both PT1 and PT7 have nuclease activity. To test our hypothesis, we assayed the nuclease activity of purified PT1 and PT7 mutants against plasmid and bacterial genomic DNA. The result indicated non-specific endodeoxyribonuclease activity of PT1 and PT7 mutants (Figure 6A-B). This result demonstrates that the point mutants retain some activity of the wild-type proteins. PT7 mutants degraded genomic DNA faster than they did plasmid DNA (Figure 6B). The catalytic activity was abolished in the presence of EDTA acting as a chelator, indicating metallonuclease activity. In contrast, no nuclease activity was shown against tRNA, confirming DNase but not RNase activity (Figure S11A). Next, we tested the capability of the purified immunity proteins to neutralize the toxin’s catalytic nuclease activity. The results indicated that the catalytic activity of PT1 mutants was abolished when it was incubated with PIM1 protein, confirming the immunity activity of purified PIM1 *in vitro* (Figure 6C). Surprisingly, PIM7 could not abolish the catalytic activity of PT7 mutants *in vitro* (Figure S11B), possibly due to the nuclease assay experimental conditions or due to the mutations.

**Figure 6:**
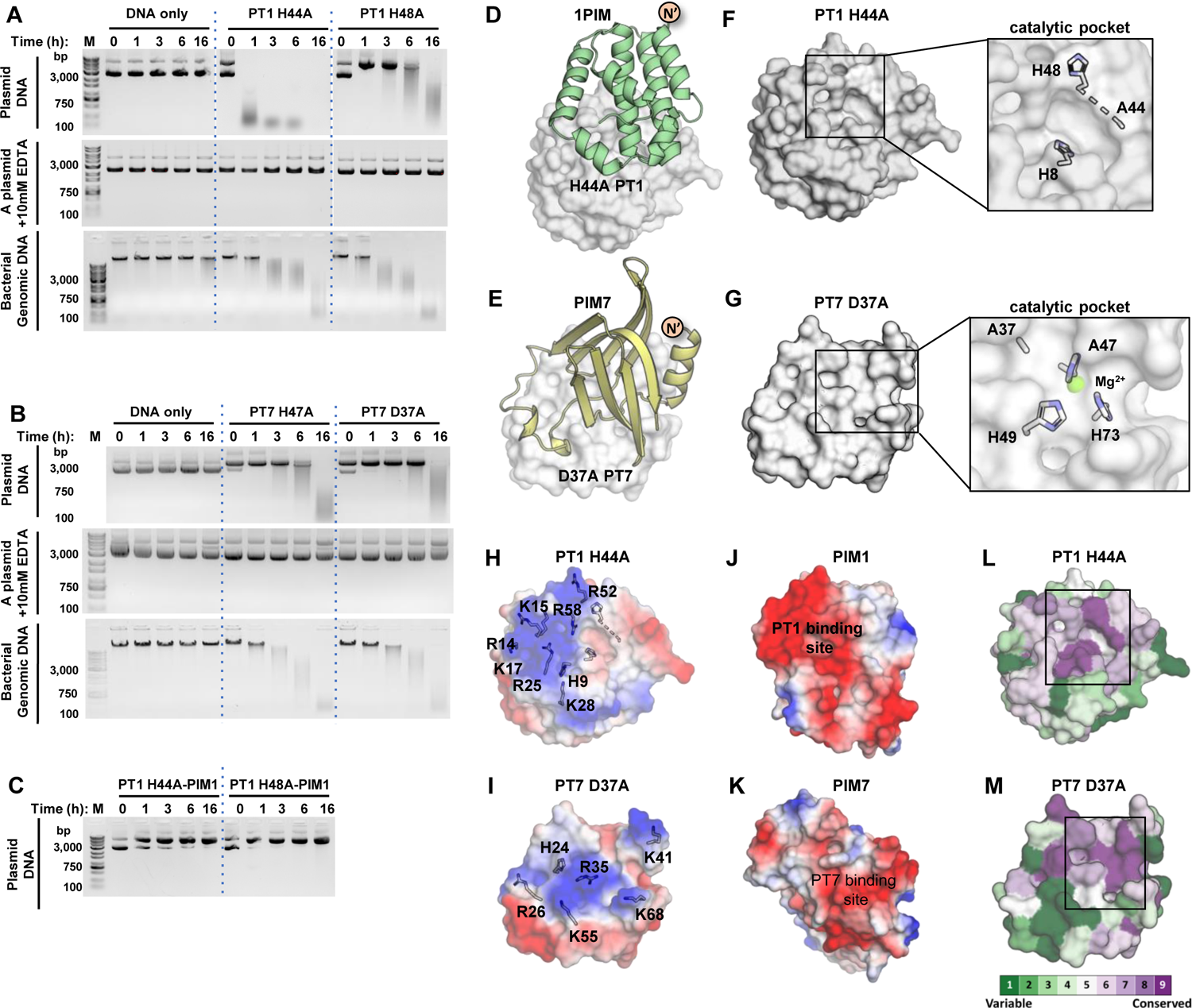
Biochemical and structural characterization of PT1 and PT7 mechanisms of action. **(A-C)** In vitro DNase activity assay of PT1 and PT7 mutants. Purified plasmid (Top and middle) or *E. coli* genomic DNA (bottom) were co-incubated with buffer (DNA only) or with purified PT1 H44A and PT1 H48A **(A)** or PT7 H47A and PT7 D37A **(B)** proteins in the presence of Mg2+ (top and bottom) or 10mM EDTA (middle) at 37 °C for different time points. The integrity of DNA was visualized on 0.8-1% agarose gel. The results indicate non-specific Endo-deoxyribonuclease (Endo-DNase) activity of PT1 and PT7. **(C)** Anti-toxin activity of PIM1. The purified plasmid was co-incubated with purified PT1 H44A-PIM1 and PT1 H48A-PIM1 complexes at 37 °C for different time points. The integrity of DNA was visualized on 1% agarose gel. The results confirm the anti-toxin activity of PIM1. **(D and E)** The crystal structure of PT1 H44A-PIM1 **(D)** and PT7 D37A-PIM7 **(E)** complexes. The PTs are shown in surface presentation and colored in gray, and PIMs in cartoon presentation. **(F and G)** The catalytic pocket of PT1 **(F)** and PT7 **(G)**. The residues that were predicted to be part of PTs catalytic sites are labeled and shown in stick presentation. The structures indicate that the catalytic residues are clustered together to form the catalytic pocket. **(H-K)** Electrostatic surface potential analysis of PT1 H44A **(H)** and PT7 D37A **(I)** indicates a positive surface in the vicinity of the catalytic site, suggesting a DNA binding site. The positive residues are labeled and shown in a stick presentation. Electrostatic surface potential analysis of PIM1 **(J)** and PIM7 **(K)** indicates that the PTs binding site is negatively charged. **(L and M)** ConSurf analysis of the PTs. Surface presentation of PT1 H44A **(L)** and PT7 D37A **(M)** with the residues colored by their conservation grades (1 to 9, where 1 is the most variable and 9 is the most conserved) indicates the high conservation of the catalytic site residues. The catalytic site is marked by a rectangle.

To investigate the structural basis for PT1 and PT7 catalytic activity, we solved the crystal structure of PT1 H44A-PIM1 and PT7 D37A-PIM7 complexes (Figures 6D-E, and Table S10). The structures confirm the new protein topology of PT1 and PT7 that do not resemble the topology and structure of any known protein (Figures S11C-F). Structural analysis of the PT proteins confirmed that the residues predicted to have a high propensity to serve in PT1 and PT7 catalytic sites are clustered together to form a putative catalytic pocket (Figures 6F-G). For PT7, an indicative electron density was observed that could be fitted by a metal ion (Mg^2+^, Figure 6G). Electrostatic potential surface analysis of PT1 and PT7 indicated a positive surface near the catalytic sites, suggestive of a binding site for the negatively charged DNA (Figures 6H-I).

Structural analysis of the PT-PIM complexes indicated a buried surface area of 1096 Å^2^ for the PT1-PIM1 complex corresponding to 17.9% of PT1 and 15% of PIM1 total surface area and 907Å^2^ for the PT7-PIM7 complex corresponding to 12.6% of PT7 and 17.2% of PIM7 total surface area. In both complexes, the PIMs hinder the catalytic pocket (Figures 6D-E), and the PT binding sites are negatively charged (Figures 6J-K), similarly to the negatively charged phosphate backbone of a DNA molecule. Analysis of the interactions between the PTs and PIMs indicated dominancy of polar interactions (hydrogen bond and salt bridge, Figure S11G). For the PT1-PIM1 complex, most salt bridge interactions are formed between the negatively charged residues surrounding the catalytic pocket. In contrast, for the PT7-PIM7 complex, most of the salt bridge interactions are formed between the catalytic residues and PIM7 D17, located at the tip of the α1-βA loop (Figure S10H).

Analyzing the conservation of the surface residues of the PT1s and PT7s toxins using the ConSurf server ^49^ indicated the high conservation of the catalytic residues (Figures 6L-M and S11I-J). Similarly, ConSurf analysis of the PIM1 proteins revealed high conservation of the PT1 binding residues (Figure S11L). In contrast, analysis of the PIM7 proteins indicated only partial conservation of the PT7 binding residues (Figure S11M), suggesting that the binding site undergoes rapid evolution.

## Discussion

Polymorphic toxins are multi-domain proteins involved primarily in inter-bacterial competition that were discovered nearly a decade ago ^15^. In recent years it is becoming clear that polymorphic toxins play important roles in various bacteria, archaea and temperate phages, and their toxins can serve as DNAses, RNAses, rRNA ADP-ribosylating and membrane depolarization enzymes, metallopeptidase, and cell wall biosynthesis inhibitors ^9–11, 14, 16, 19, 21, 50–56^. Here, we developed a new algorithm to predict novel toxin domains (PTs) located at the C-terminus of polymorphic toxin proteins across over 100,000 bacterial genomes. We also predicted the cognate immunity genes (PIMs) of the new PTs. This *in silico* analysis was experimentally confirmed in heterologous expression assays in *E. coli* and *S. cerevisiae* and led to discovery of nine classes of potent PTs that are toxic to bacteria or yeast and five cognate PIMs that rescue cells from toxin activities. Five predicted toxins failed to show toxicity in our systematic experimental pipeline (Table S3). It is plausible that some of our false negative predictions kill bacterial taxa that we have not tested such as Gram-positive bacteria which are well-known targets of polymorphic toxins ^10, 50, 55, 56^. Thus, expanding the host range for heterologous expression in other model organisms might yield favorable results in future studies.

The novel PTs are part of various polymorphic toxins that are based on the N-terminal trafficking domains genetically associated with various microbial secretion systems, such as T5SS, T6SS, T7SS, and eCIS (Figure 1B). Future work should establish a biochemical association between the PT-encoding effectors and their putative secretion systems. Interestingly, the short PT-encoding genes we experimentally validated as toxins are by definition single domain polymorphic toxins. This finding may suggest that these specific genes either function as cargo effectors of different secrtion systems which are likely loaded via adaptor proteins ^57–59^, or function in self groth inhibition similarly to classical toxin-antitoxin systems in phage defense^60, 61^.

We analyzed the ecological and phylogenetic properties of the PTs and identified that they are encoded by 2.2% of all sequenced microbes. We provided a proof of concept that the new PTs can quickly and efficiently target different bacterial and fungal cells from the cell exterior and thereby can be used as alternative antimicrobial agents against antibiotic resistant bacteria and fungi. Moreover, the fact that the PT-GFP fused proteins can penetrate fungal cell membranes is intriguing due to their large size. Future studies should identify the mechanism by which these proteins are imported and will investigate if this effect is dependent on other molecules present in the lysate of PT-expressing cells. It will be also interesting to characterize the PTs’ target organism range and whether they evolved primarily to target a specific group of organisms.

We shed light on the versatile mechanism of action of the toxin through imaging of PT expressing cells using fluorescence microscopy. We can indicate potential cellular targets of toxin activity such as DNA, cell wall, cell membrane damaging, or the cell division process. We then predicted the protein folding of the novel toxins and experimentally confirmed critical residues, which are often buried in predicted enzymatic active sites, that are likely participating in substrate catalysis. Additionally, we solved the structure and mode of actions of two novel nucleases: mutants of PT1 and PT7, and their cognate PIMs. The mutants were non-toxic *in vivo* (Figure 3B) but retained their biochemical activity *in vitro* (Figure 6A-B), suggesting that the target cells can overcome the mutant activity through DNA repair mechanisms. Biochemical and structural data revealed that PT1 and PT7 act as efficient DNAses and their PIMs occlude the enzymes’ active sites. Although PT1 and PT7 are nucleases of novel fold they share their positively charged DNA binding domain with proteins of similar function ^62–66^. Future study needs to be done to decipher the structure-function of the other PTs and might be valuable for developing effective approaches to combat bacterial and fungal infections.

Overall, this work significantly expands the space of toxin domains of PTs and uncovers in the macro-scale the broad role of these toxins through their correlation with different phylogenetic and ecological data and potential target organisms, and in the micro-scale the PT cellular function within the attacked cells. Using our approach we confirmed that the modular nature of polymorphic toxin proteins serves as a good predictor for toxin domains. We strongly suggest that additional PT domains and cognate PIMs remain hidden in the *Terra Incognita* of microbial genomes and some fine-tuning of the parameters we used in computational prediction can expand these gene functions much further.

## Materials and Methods

### Collection of known domains found in polymorphic toxins

We collected a list of Pfam domains ^25^ and one TIGRFAM domain that are common in polymorphic toxin proteins. These served as anchors for our search for toxin C-terminal domain. The list included trafficking domains, repeat domains, pre-toxins, and toxins. Trafficking domains included: VgrG, PAAR ^3^, DUF4150, HCP, LXG ^10^, DUF4157 ^28, 54^, MafB ^67^, Wxg100_esat6, Phage_min_cap2 ^15^, SpvB ^15^, PrsW-protease, DUF4280 ^3^, and TANFOR ^19^. Repeat domains included: Haemagg_act ^67^, filamentous hemagglutinin repeats ^67^, RHS, RHS repeat ^15^, TcdB_toxin_midN, DUF637, and ALF. Pre-toxins included PT-HINT ^15^, PT-TG ^15, 67^, and PT-VENN. Toxins domains included: RNAse_A_bac, AHH, Ntox21, Ntox27, Colicin_D, Ribonuclease, XOO_2897-deam, SCP1201-deam, HNH, Tox-REase-7. We then expanded this list iteratively using the following procedure, each time treating the newly expanded list as the input for the procedure, until the list was no longer expanded (Figure S12). For each known PT domain (‘anchor domain’), we iterated over the Pfam architectures that contain it ^25^ and examined the domains in each architecture (namely, an annotated protein sequence) that are yet unknown PT domains (denoted as ‘potential PT domain’). If a potential domain in some architecture containing some anchor domain met one of the conditions according to the schema in Supplementary Figure S10, it was added to the expanded list with an annotation of a certain type (trafficking, repeat, pre-toxin, toxin). Assessment of domain type was done using its Pfam name and description. Each type was assigned with some name-keyword and description-keyword that were manually extracted from Pfam description of known PT domains (e.g. “tox” as the name-keyword for toxin domains, and “bind” for trafficking). Each presence in the name of some name-keyword contributed 1.5 to the score, and each presence in the description of some description keyword contributed 0.5. If name or description contained the string “polymorphic toxins” it contributed 1.5 to all PT domain types. If the PT domain type with maximum score had a score equal or higher than 1.5 the domain was assigned with this type, otherwise it was annotated as a domain whose type could not be assessed and it was discarded. Putative domains that their assessed type didn’t make sense were omitted manually. The final list of 217 domains is listed in Table S1.

### Obtaining initial toxin domain candidates

We used 105,438 microbial genomes that were downloaded from the IMG database in 2018 ^68^. We first defined candidate polymorphic toxin genes. Namely, those genes that contain multiple domains found in polymorphic toxins. Gene domain annotations were obtained from IMG data. We calculated that the median domain size in pfam is 133 aa (as of 2020) and therefore we filtered in only genes encoding proteins of at least 2.5 domains (133 aa x 2.5) that contained at least one domain that is a known\putative trafficking, repeat or pre-toxin domain according to the expanded list from the previous analysis. We defined the C-terminus as the last 133 amino acids of the protein. The initial list included 2,633,634 C-terminal sequences from 82,253 genomes. These were clustered using cd-hit version 4.8.1 (40% identity) ^69^ to obtain 152,150 clusters. These clusters represent unique C-terminal domains that are candidate toxins from polymorphic toxins. However, we were aware that this list included many false positive predictions and in the next steps we worked to increase the signal to noise ratio in predictions.

### Filtering putative toxin domains for taxonomic diversity

We took each candidate’s cluster and obtained the genus of each member from the IMG genome metadata. We omitted candidates which had a total of less than three unique genera among their cluster members. This left us with a total of 11,196 candidates. The purpose of this step was to maintain only domains that are more likely to be functional and therefore selected during microbial evolution or horizontally transferred between multiple genera.

### Filtering putative toxin domains for novelty

To represent the sequence space of each cluster we calculated the multiple sequence alignment of the cluster sequences with Clustal Omega version 1.2.4 ^70^ and then, using the MSA, we constructed a profile hidden markov model (HMM) using HMMER’s ^71^ hmmbuild program. We used the hmmsearch program to search all of the candidate hmms against the Pfam-A fasta file (e-value <= 0.001). Each candidate with some hit to a known domain was omitted, to give us a total of 1,986 candidates that had no Pfam hits. We performed a more sensitive novelty check against additional databases only for top-scored candidates (see below: “Re-filtering toxins for novelty using a more sensitive sequence-based search”).

### Scoring putative toxin domains for modularity

We used the modularity feature of polymorphic toxins to verify that the novel domains appear in various architectures of polymorphic toxins, thereby supporting their polymorphism. Namely, it is more likely that a real toxin domain (PT) will be fused to multiple polymorphic trafficking and repeat domains that can therefore be secreted by various secretion systems. For each cluster, we searched with DIAMOND ^72^ (e-value <= 0.001) the cluster representative sequence (decided by cd-hit) against NCBI nr, taking only the subject proteins that had a hit at their C-terminal (alignments that end maximum of 103 aa from the C-terminus), as we were interested in proteins that manifest the polymorphic toxins template architecture. Then, we annotated all these protein domains using hmmscan ^71, 72^ (e-value <= 0.001) against Pfam and created the domain architecture for each of the proteins. For each architecture, we kept a domain annotation only if it was in the expanded list of PT domains, and only if it wasn’t at the C-terminus (ended at least 103 aa away from the C-terminus). Additionally, if a given architecture had consecutive occurrences of a repetitive domain, we treated it as a single domain occurrence. Finally, we counted all unique architectures, and defined this number as the modularity score of the cluster (namely, the candidate toxin). We treated all candidates with a modularity score of at least four as top-scored candidates. We manually omitted candidates that were only derived from proteins of many different combinations of some repetitive domains, without trafficking or pre-toxin domains for instance.

### Re-filtering toxins for novelty using a more sensitive sequence-based search

We took the multiple sequence alignment of each top-scoring candidate and validated their novelty with HHpred server ^73^ against PDB, SMART, NCBI CDD, Pfam-A, omitting candidates with significant hits (probability > 95%, e-value <= 0.001). Additionally, we searched each candidate’s profile hmm with hmmsearch against Swiss-Prot, omitting every candidate with a significant hit of a toxin or an enzyme (e-value <= 0.001, query cover >= 50%).

### Selecting genes for synthesis

One of the challenges we faced was to identify the toxin domain borders, so we can test our computational predictions in molecular biology experiments. The strategy we employed was to select single-domain protein-coding genes that contain the candidate toxins domains. For each candidate, we searched its cluster representative sequence using BLASTP ^74^ (e-value <= 0.01) against nr and DIAMOND ^72^ (e-value <= 0.01, ‘very sensitive’ option) against nr and IMG databases and manually extracted sequences of significant hits that had long alignment coverage of the subject sequence and did not have significant hits when searched in HHpred (against Pfam-A, NCBI CDD, SMART, PDB databases, probability > 95%, subject cover >= 50%). Seven of the final chosen sequences had length that was abundant among the hits of the cluster sequences. Here we assumed that a high frequency of specific gene length will be the most functional version of our candidate toxin due to the length conservation.

### Obtaining candidates for immunity genes

For seven of our final candidates we predicted an immunity protein as well by taking the gene downstream to the single domain PT gene in the genome, and manually validating that it was in synteny with the single domain PT gene.

### Architectures analysis

Novel toxin sequences were searched against IMG with DIAMOND (very sensitive, evalue <= 0.001, query cover >= 80%). For each toxin, we analyzed the domain architecture of the proteins that had hit of the toxin using hmmscan (c-e-value <= 0.001, query cover >= 50%) against a profile HMM database containing Pfam and TANFOR domain ^75^ for TIGRFAM ^76^ (Table S4). We considered an overlap of domain annotation to be an overlap of >= 50 amino acids of two domain annotations, and dealt with it by keeping the annotation with the lower c-e-value. Hits of validated immunity proteins were found using DIAMOND as above. We extracted all hits of immunity proteins which were immediately downstream of some occurrence of their matching validated toxin using IMG gene IDs, and showed it in Figure 1B as well.

### Phylogenetic analysis

In order to build a phylogenetic tree that spanned our bacterial genomes database, we chose a representative genome from each taxonomic Family as the resolution for our dataset, which yielded a tree in which each leaf represents a taxonomic family. As a basis for comparison, we used universal marker genes. Specifically, we used 29 COGs out of 102 COGs that correspond to ribosomal proteins ^77^. We aligned each COG from each representative genome using Clustal Omega version 1.2.4 ^70^. We then concatenated all the COGs alignments for each genome, filling missing COGs with gaps. We used this 29 marker gene concatenated alignment as input to FastTree2 with default parameters ^78^ to create the phylogenetic tree. To exhibit the distribution of occurrences of novel toxins, we first identified orthologs of the tested genes by searching for them in bacterial genomes in IMG using DIAMOND with query cover >=80%, e-value <= 0.001 and ‘very sensitive’ option. R’s ggtree v2.4.2 and ggtreeExtra v1.0.4 packages ^79, 80^ were used to plot the tree as well as the percentage of genomes in the family having at least one occurrence of the toxin according to our DIAMOND homology results. We present in the tree phylum annotation only for families that encode at least one of the novel toxins, or families that belong to a phylum that contains more than two families according to our tree.

### Calculation of percentage of sequenced bacteria encoding the novel toxins

Homologs of novel toxins were obtained from the IMG microbial genome database that was downloaded in 2018 using DIAMOND ^72^ with the following parameters: e-value <= 0.001, query-cover >= 80%, ‘very sensitive’ option). The percentage of sequenced bacteria encoding the novel toxins was calculated based on the number of unique genomes encoding these homologs (n=1339) and total number of IMG genomes with ‘Bacteria’ value in the ‘Domain’ field according to the IMG metadata (n=62075).

### Ecosystem enrichment analysis

The enrichment analysis assigns an odds ratio that quantifies the enrichment of metadata labels using a Fisher exact test. To correct for taxonomic bias, a population-aware enrichment analysis based on Scoary ^81^ version 1.6.16 was used. The inputs to Scoary were (1) a guide tree, (2) a presence-absence file based on occurrences of toxins (see phylogenetic analysis), indicating if a genome encodes for the toxin, (3) metadata files that indicate for each genome if they possess a certain characteristic, e.g. whether it was isolated from soil. One guide tree was created per metadata category, based on universal marker genes (see phylogenetic analysis) and only those entries with at least 25 instances in the database were included. Statistical significance for both the naive Fisher exact test and for the phylogeny-aware Scoary test were calculated, displayed as -log10 of the q-values. Correlations with q-value <= 0.05 for both Fisher exact test and Scoary test were considered significant. q-values (False discovery Rates) were obtained using the Benjamini-Hochberg procedure ^82^.

### Conserved amino acids analysis

For each toxin we obtained hits in IMG database using DIAMOND (e-value <= 0.001, query-cover >=80%, ‘very sensitive’ settings). For PT2, PT8 we had a small number of hits so we used DIAMOND to search the obtained hits again in the IMG database and used these hits. For each toxin we clustered the hit alignments using cd-hit version 4.8.1 (90% identity) and calculated the MSA of the clusters’ representatives using Clustal Omega version 1.2.4. We then used each of these MSAs to generate HMM logo for each toxin with skylign.org ^83^ using the following parameters: Create HMM - keep all columns, Alignment sequences are full length, Information Content – All. The same logos, but generated with ‘Create HMM - remove mostly-empty columns’ are shown in Figure S5. Additionally, we scored each position using Jensen-Shannon Divergence ^43^. Based on this information, we manually chose each toxin top-scoring positions with amino acids that have high propensity to be active in catalysis in enzymes ^44^.

### Structural analysis

Predicted structures of the experimentally validated toxins were obtained using RobbeTTAFold ^39, 84^. Generated 3D models were visualized using ChimeraX ^85^. In order to demonstrate co-localization in the microenvironment of catalytic amino acids, we chose cartoon mode and highlighted specific residues (without their hydrogens). Visualisation of the surface with coloring of chosen residues was used to demonstrate catalytic pockets. We used Dali server ^86^ in order to find proteins with similar structure, and used the ‘Matches against PDB25’ results.

### Bacterial strains and strain construction

Candidate toxin and immunity gene sequences were retrieved from IMG, synthesized (codon optimized for *E. coli*), and cloned into either pBAD24 (Thermo Fisher Scientific, 43001) or pET28a plasmids by Twist Bioscience, respectively. Growth control for the experiments was an empty vector. The plasmids were then transformed using the previously described TSS method ^87^ into *E. coli* BL21 (DE3) strain. For Toxin-immunity protection assay, *E. coli* BL21 (DE3) cells harboring pBAD24 plasmids with toxic genes were co-transformed with pET28 vectors harboring cognate immunity genes or with empty vectors as control (Table S2, S12).

### Site-directed mutagenesis and PT-*gfp* constructs

Chosen residues were substituted with alanine using Q5® Site-Directed Mutagenesis Kit (NEB) according to supplier protocol. The mutante genes and PT-*gfp* were constructed in pBAD. In brief, primers for PCR were generated using automated software offered by NEB. pBAD vectors harboring the genes were linearized using the primers in inverse PCR and then incubated in 1X KLD mix provided with the kit for 5-10 minutes. The mixture was transformed into *E. coli* DH5α strain (NEB), screened by colony PCR, and validated by Sanger sequencing. Plasmids from positive clones were prepared using a mini prep kit and re-transformed into *E. coli* BL21 (DE3) strain. Residues that failed this method were substituted by re-cloning the gene as two overlapping fragments using NEBuilder® HiFi DNA Assembly Cloning Kit (NEB) according to supplier protocol and recommendations. In brief, target genes were PCR amplified as two overlapping fragments having the point of mutation designed in the midst of the overlap. pBAD24 vector linearized with NcoI was incubated with fragment and NEbuilder enzyme mix for 30-45 minutes. The mix was cloned into DH5α strain and validated as mentioned above (Tables S10-S11).

### Drop assays and Growth curves

For bacterial drop assays and growth curves, overnight cultures of the strains harboring the vectors were grown in LB supplemented with ampicillin, 100 mg/liter (Tivan biotech) or also with kanamycin 50 mg/ml (Tivan biotech) for toxin-immunity cells with shaking (200 rpm). The cells were grown on LB plate or in LB liquid with ampicillin (100mg/ml) and 0.2% arabinose or 1% glucose for cells containing toxin or mutant toxin only. For toxin-immunity experiments, ampicillin (100mg/ml), kanamycin (50 mg/ml), 0.2% arabinose and 0.01mM IPTG respectively. For drop assay, cultures were normalized to OD600 = 0.3 and then serially diluted by a factor of ten. Dilutions were spotted as three biological replicates on LB agar containing as were described above. The plates were incubated overnight at 37 °C. Results were documented using amersham ImageQuant 800. To construct growth curves, cultures were normalized to OD600 = 0.03 then incubated at 37 °C with shaking at 150 rpm for up to 12 h in a microplate reader (Synergy H1, BioTeck). OD600 were measured continuously over 12 h. Data analysis was performed with R (version 4.0.5) software, ‘ggplot2’ and ‘growthrates’ packages ^88, 89^.

### Yeast gene synthesis, heterologous expression, and drop assay, and growth curves

Heterologous expression of putative toxin in yeast candidate sequences were retrieved from IMG. They were synthesized, following codon adaptation to yeast by Twist Bioscience and were cloned into pESC -leu galactose inducible plasmids. The plasmids were then transformed into *Saccharomyces Cerevisiae* BY4742 strain (Table S12). Overnight cultures of the strains harboring the vectors of interest were grown in SD-leu media. The cultures OD was normalized OD600 = 0.3, washed once with water and then serially diluted by a factor of ten. Dilutions were spotted as three biological replicates on SD-leu agar containing 2% glucose or 2% galactose and plates were incubated for 48 hours at 30°C. Results were documented using amersham ImageQuant 800. To construct growth curves cultures were normalized to OD600 = 0.03 then incubated at 30 °C with shaking at 150 rpm for up to 40 h in a microplate reader (Synergy H1, BioTeck). OD600 were measured continuously over 40 h. Data analysis was performed with R (version 4.0.5) software, ‘ggplot2’ and ‘growthrates’ packages ^88, 89^.

### Statistical analysis

In each drop assay experiment, CFU measurements were compared between the induction of the toxin alone and and between all other groups. An unpaired t-test was used to test the hypothesis that the means are different using Scipy 1.9.1 python package, followed by FDR correction for multiple comparison. *P* value <0.05 was considered significant. Plots were generated using ggplot2 package in R.

### Fluorescence microscopy for *E.coli* cells

Microscopy of induced cells *E. coli* BL21 cells that contain the wild-type PTs, the mutated PTs or PT-PIM pairs were grown in LB media at 37 °C. When growth reached an OD600 of 0.3, bacteria were induced with 0.2% arabinose for PTs or 0.2% arabinose and 0.01 Mm IPTG for PT-PIM’s strains for 40 minutes. 300 μl of the induced samples were centrifuged at 8,000xg for 2 minutes at 25 °C and resuspended in 5 μl of Dulbecco’s phosphate-buffered saline (DPBS) (Thermo Fisher Scientific 14200075), supplemented with 1 mg/ml membrane stain FM1-43 for bacteria and FM4-64 for yeast (Thermo Fisher Scientific T35356 and T13320) and 2 μg/ml DNA stain 4,6-diamidino2-phenylindole (DAPI) (Sigma-Aldrich D9542-5MG). Cells were visualized and photographed using an Axioplan2 microscope (ZEISS) equipped with an ORCA Flash 4.0 camera (HAMAMATSU). System control and image processing were carried out using Zen software version 3.1 (Zeiss).

### Liposome preparation (LUVs) and size characterization and Calcein-release assay

*E. coli* extract (100500P, Sigma) was dissolved in chloroform and the organic solvent was evaporated using the first nitrogen stream, followed by 3 h of vacuum pumping. The lipid film was then hydrated with 3 mL calcein solution (70 mM, osmol=300 mOsmo/kg) at 35°C. The resulting MLV suspensions were sonicated for 15 min at 35°C to disperse larger aggregates. The vesicles were subsequently downsized by extrusion (Lipex, Northren Lipids Inc) through 400 nm, 200 nm and 100 nm polycarbonate membranes. The extrusion was performed 11 times through each membrane at 35°C. Liposomes were then separated from an excess of free calcein by 48 h - dialysis (Float-A-Lyzers, Sigma-Aldrich) against PBS. The size of the liposomes was measured with a ZetaSizer Nano ZS (Malvern Instruments, UK) at 25°C. Triplicate measurements with a minimum of 10 runs were performed for each sample.

The effect of the PT on pore formation in the lipid membrane was monitored by calcein dequenching methods as described elsewhere ^90^. The calcein-loaded liposomes (25 µL at concentration ∼1 mM) were added into 50 µL of cell lysates of PT-expressing *E. coli* cells. Continuous monitoring of calcein fluorescence (excitation 470 nm, emission 509 nm) was done at interval of 20 min for a period of 24 h (shaking 100 rpm) at 37 °C, using a ClarioStar microplate reader (BMG LABTECH GmbH, Germany). The final fluorescence intensity, which represents maximal fluorescence of free calcein was determined following the solubilization of vesicles with Triton X-100 (1% v/v) and controlled with the value of calcein obtained from solubilized liposomes in the peptide-free medium. All statistical assays performed were analyzed using Analysis of Variance (ANOVA) and then Tukey’s test using OriginJ software v1.52i (NIH, USA). P-values <0.05 were considered statistically significant.

### Preparation of PT-expressing cell lysates

Overnight cultures of *E. coli* expressing PTs were diluted 1:100 in 100 ml LB medium and grown at 37°C (250 rpm) for 1 h 45 min. The expression of each PT was induced by the addition of arabinose (final concentration 0.02%) and cells were further incubated at 37°C (250 rpm) for 3h. Cells were then centrifuged at 4,000 rpm for 20 min at 4°C and samples kept on ice throughout the cell lysates preparation. Pellets were resuspended in a 10 ml Lysate buffer containing 50 mM Tris-HCl (pH 7.4) and 150mM NaCl. The resuspended cells were then sonicated (10 pulses, 10 sec and cell lysates were then centrifuged at 10,000*g* for 30 min at 4°C and the supernatant was divided to 1 ml tubes and was kept in −20c prior to antibacterial and antifungal assays.

### Antibacterial assays

For growth curves, overnight cultures of *B. Subtilis* or *E.coli* were grown in LB with 1:50 lysates of *E.coli* PT expressing cells. Cultures were normalized to OD600 = 0.03 then incubated at 37 °C with shaking at 150 rpm for up to 12 h in a microplate reader (Synergy H1, BioTeck). OD600 were measured continuously over 12 h. Data analysis was performed with R (version 4.0.5) software, ‘ggplot2’ and ‘growthrates’ packages ^88, 89^. To test Propidium iodide (PI) staining of *B. Subtilis*, *B. Subtilis* cells at mid logarithmic phase were incubated with lysates of *E.coli* PT expressing cells for 2 hours, then the cells were harvested and washed twice in PBS. Cells were treated with 5 μg/ml PI for 2 min and were visualized by fluorescent microscopy (see above).

### Antifungal assays

Growth curves were determined by the microdilution method recommended by Clinical and Laboratory Standard Institute (CLSI) M38-A2 and M27-A2 for filamentous fungi and yeasts, respectively. The experiments were performed in 96-well plates using GMM-media (1% [wt/vol] glucose, 1 × original high-nitrate salts, trace elements, pH 6.5; trace elements, vitamins, and nitrate salt compositions were as described previously ^91^) for *A. fumigatus* and *A. nidulans*, and Saboraud Dextrose (HiMedia M063) for *F. oxysporum*, *C. albicans*, *C. auris* and *C. neoformans*. Each well was filled with 175 μL of serial broth dilutions containing different concentrations of the bacterial lysates tested and inoculated with 5 μL of a fresh culture of each organism (10^5^ CFU or conidia mL−1 ^92^). 20 μL of Resazurin sodium salt (Sigma Aldrich, R7017) was added into the above wells as a colorimetric indicator of metabolic activity. The plates were incubated for 48 h at 37°C for *A. fumigatus*, *A. nidulans* and *C. auris*, and 48 h at 28°C for *F. oxysporium*, *C. albicans* and *C. neoformans*. Fluorescence intensities (λ = 590 nm) of treated and untreated samples for each strain were obtained using a Spectra Max SpectraMax i3 (Molecular Devices) after 48 h of incubation and used to calculate the percentage of growth, according to the manufacturer’s protocol. The percentage of toxicity was calculated with the following equation, using absorbance: % toxicity = (100 x (No lysate - sample incubated with PT)).

Germination rate and hyphae sizes were performed according to Rocha et. al 2015 ^93^, with some modifications. To determine the germination rate of conidia in the presence of the tested PTs, 1×10^4^ conidia of *F. oxysporum* were inoculated onto 8-well glass coverslips containing 200 µL of RPMI 1640 medium (Thermo Fisher Scientific) and 25% (v/v) of lysates of *E.coli* PT expressing cells, incubating at 28°C for 6 hours, and then photographed. A conidiospore was counted as germinated if it possessed a germ tube and the germ tube length was at least as long as the spore. Hyphae size was determined microscopically by measuring 100 microconidia in 10 different areas. After 12 h of incubation, the size of the hypha was measured, and it was possible to observe the inhibition in the presence of PTs. Statistical analysis was performed using a one-way ANOVA with Dunnett’s post hoc test compared to the control condition. Cells were visualized and photographed using a confocal laser scanning fluorescence microscope LSM 900 with Airyscan 2 (ZEISS). All images were acquired using a 63X, oil immersion objective. Images were recorded in brightfield mode and in confocal mode using 488 excitation and laser channels. System control and image processing were carried out using Zen software version 3.1 (Zeiss).

### Protein expression and purification for biochemical and structural studies

PT1 and PT7 mutants were cloned into the pET28 vector as His-tagged proteins with a TEV cleavage site downstream to the His tag (His-TEV-PT). The expression vector was transformed into BL21(DE3) strain and cells were inducted by 1mM IPTG overnight at 30 °C. On the next day cells were collected by centrifugation, resuspended with 100 mM NaCl, 5mM MgCl_2_, 0.2% Triton X-100, 20 mM Tris-HCl pH 7.5 buffer supplemented with DNase, Lysozyme and protease inhibitors. Cells were mechanically disrupted using Cell Disruptor, and the lysate was centrifuged for 30 min at 4 °C. The supernatant, containing the soluble proteins was loaded on HisTrap HP column (Cytiva) washed in 500 mM NaCl, 20 mM Tris-HCl pH 7.5 buffer and eluted using 100 mM NaCl, 20 mM Tris-HCl pH 7.5 and 250mM Imidazole buffer. The eluted protein was then purified by size exclusion chromatography (SEC) using a Superdex-75 column (Pharmacia) in 100 mM NaCl, 20 mM Tris-HCl pH 7.5 buffer. The cytosolic expression of PT1 and PT7 point mutants resulted in a high yield expression of 25-50 mg/L.

PIM1 and PIM7 were cloned into the pET28 vector as untagged proteins and expressed in a similar protocol as the PTs. For PT-PIM complexes purification, cell lysates from PT and PIM expression were incubated for 1-2 hours at 4 °C to form PT-PIM complex. The complexes were purified on HisTrap HP column followed by SEC in a similar protocol as the PTs.

### Nuclease assay

0.5mg of plasmid DNA (purified pAdvantage plasmid, Promega), 1mg of bacterial genomic DNA (isolated from E. coli cells using genomic DNA kit, Qiagen), or tRNA (from E. coli MRE 600, Sigma) was incubated with PT mutants, PT-PIM complexes (final nuclease concentration of 5mM), or water (as control) in reaction buffer containing 10mM MgCl_2_, 100mg/ml BSA, 10mM Bis Tris propane pH 7 at 37 °C for different time points. To test metallonuclease activity we used a buffer containing 10mM EDTA instead of MgCl_2_. Nuclease reactions were analyzed on 0.8-1% agarose gel and stained with ethidium bromide.

### Crystallization and structural determination of PT-PIM complexes

Crystallization experiments were performed using the vapor diffusion sitting drop method at 20°C. Crystallization screen plates were set up by the Mosquito crystallization system (SPT Labtech) using the JCSG-IV screens (Molecular Dimension) and protein concentration of 10-15 mg/ml. Crystals of PT1 H44A - PIM1 complex were obtained using a reservoir solution of 10% PEG-8000, 0.2 M Ca-acetate, Imidazole pH 8.0. Crystals of PT7 D37A - PIM7 complex were obtained using a reservoir solution of 12% PEG-20000, 0.1 M MES pH 6.5. Prior to data collection crystals were cryoprotected with 10-15% ethylene glycol and flash-cooled in liquid nitrogen. Diffraction datasets were collected at the European Synchrotron Radiation Facility (ESRF) beamline ID-30A (MASSIF-1), Grenoble, France, at 100 K. The structures were solved by molecular replacement using Phaser ^94^ with the computational model structures as search models. Structure refinement and model building were conducted in Phenix ^95^, and model building was performed with COOT ^96^. Final refinement statistics are summarized in Table S9.

## Acknowledgements

AL is generously supported by the Israeli Science Foundation (Grants #1535/20, #3300/20, #3062/20), Alon Fellowship of the Israeli council of higher education, The Hebrew University - University of Illinois Urbana-Champaign seed grant, the Israeli Ministry of Agriculture (Grant 12-12-0002), and ICA in Israel. NN is supported by the Israeli Science Foundation (Grant #1535/20) and a PhD scholarship from Faculty of Agriculture, Food, and Environment, the Hebrew University of Jerusalem. NT is supported by the Israeli Science Foundation (Grants #1600/21, #1818/21). NS is supported by the Israeli Science Foundation (Grant # 1760/20), the Zuckerman STEM Leadership Program, the Ministry of Science and Technology of Israel (Grant #0001998), and BARD US-Israel Agricultural Research and Development Fund (Grant #IS-5492-22 R). MK and JK are supported by the European Research Council (Adv. Grant # 743016).

We thank Asaf Levy’s Lab members for critical reading, advice and helpful discussions throughout, and especially thank Alex Geller. NN is thankful to Hagit Almagor for her help during the COVID-19 pandemic. We thank the ESRF for the provision of synchrotron radiation facilities, and the ESRF ID-30A local contacts for their support and help during data collection.

