## Supplementary material for "Systematic Discovery of Antibacterial and Antifungal Bacterial Toxins": PT - supp info.docx

**Supplementary Information**

Content:

**Supplementary Figures**

Figure S1. Operon organization and protein domain architecture of classic polymorphic toxin proteins

Figure S2. Taxonomic distribution of the novel PTs..

Figure S3. The novel PTs efficiently kill bacteria and yeast.

Figure S4. PIMs provide defense against toxicity of cognate PTs.

Figure S5. Hidden Markov Models (HMM) logos for seven of the nine novel toxins

Figure S6. Toxins lead to cell death in various manners, and mutants abrogate their cellular toxicity

Figure S7. PTs 3D Fold prediction

Figure S8. Lyasets of PT-expressing *E. coli* cells lead to *B. subtillis* killing in various manners.

Figure S9. PTs form permeabilizes liposome membranes to elicit cell death

Figure S10. Cell lysates of PT expressing E. coli nhibit fungal growth

Figure S11. Biochemical and structural characterization of PT1 and PT7 mechanism of action

Figure S12. Schema for adding putative PT domains to the expanded list

**Supplementary Tables**

Table S1. Expanded list of known and putative PT domains.

Tabel S2. Novel PTs and PIMs identified in this study. Protein ID and existing protein annotation lay on the NCBI database.

Tabel S3. Candidates identified by the algorithm and failed experimental validation.

Table S4. Copy number of architectures

Table S5. All homologous proteins of PT1-9, PIM1, 3, 7-9.

Table S6. Bacterial pathogens that encode the novel PTs.

Table S7. Loci of PT3 homologs in *Pseudomonas aeruginosa* genomes from IMG.

Table S8. Correlation of bacteria encoding the novel toxins with habitats and bacterial life-styles.

Table S9. Top DALI hits (‘Matches against PDB25’ section in DALI results) of the nine novel toxins’ predicted structures.

Table S10. Data collection and refinement statistics for PT1 H44A – PIM1 and PT7 D37A – PIM7 complex structures.

### Table S11. PCR primers used for site-directed mutagenesis and for PTs-*gfp* fusion

Table S12. Bacterial and fungal strains

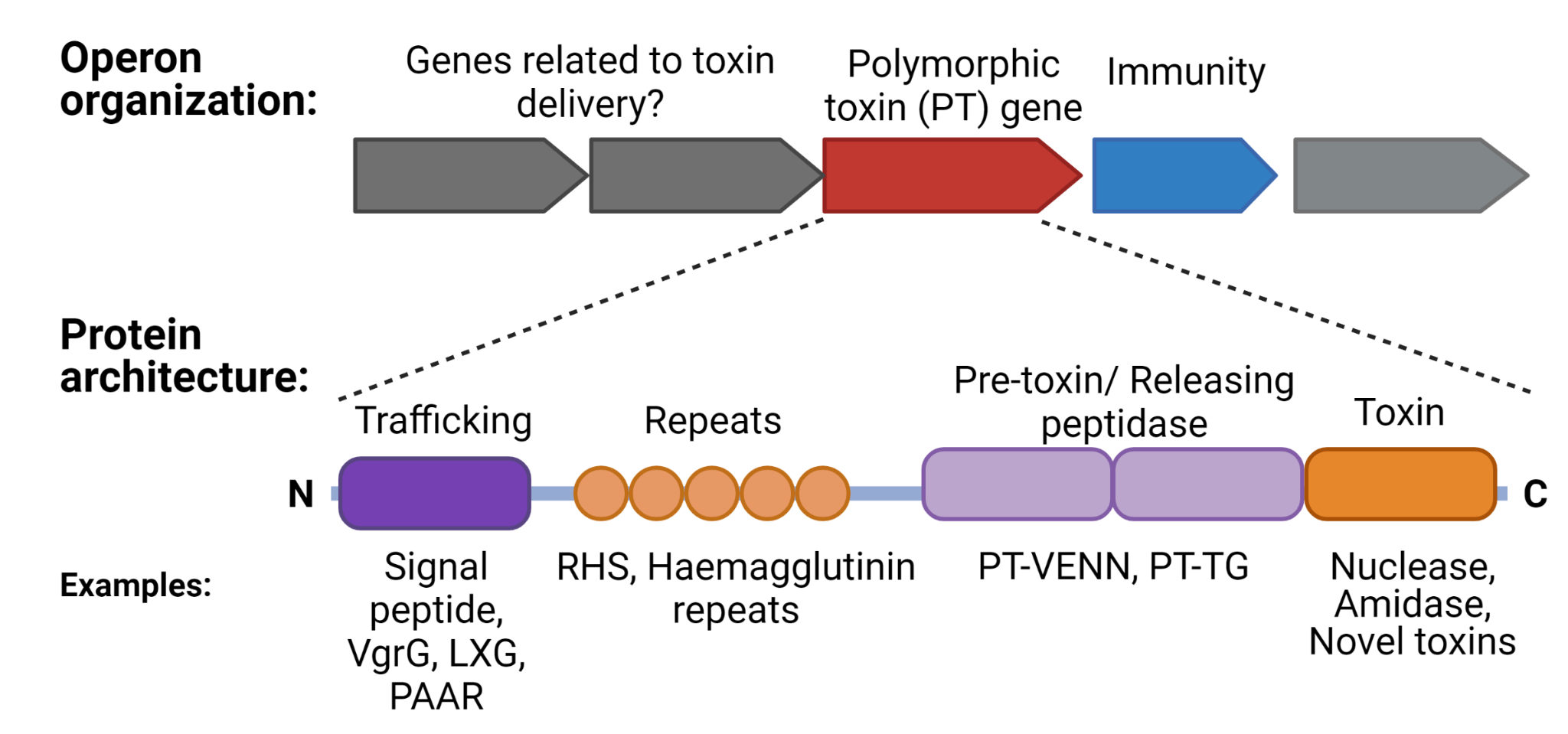

**Figure S1. Operon organization and protein domain architecture of classic polymorphic toxin proteins.**

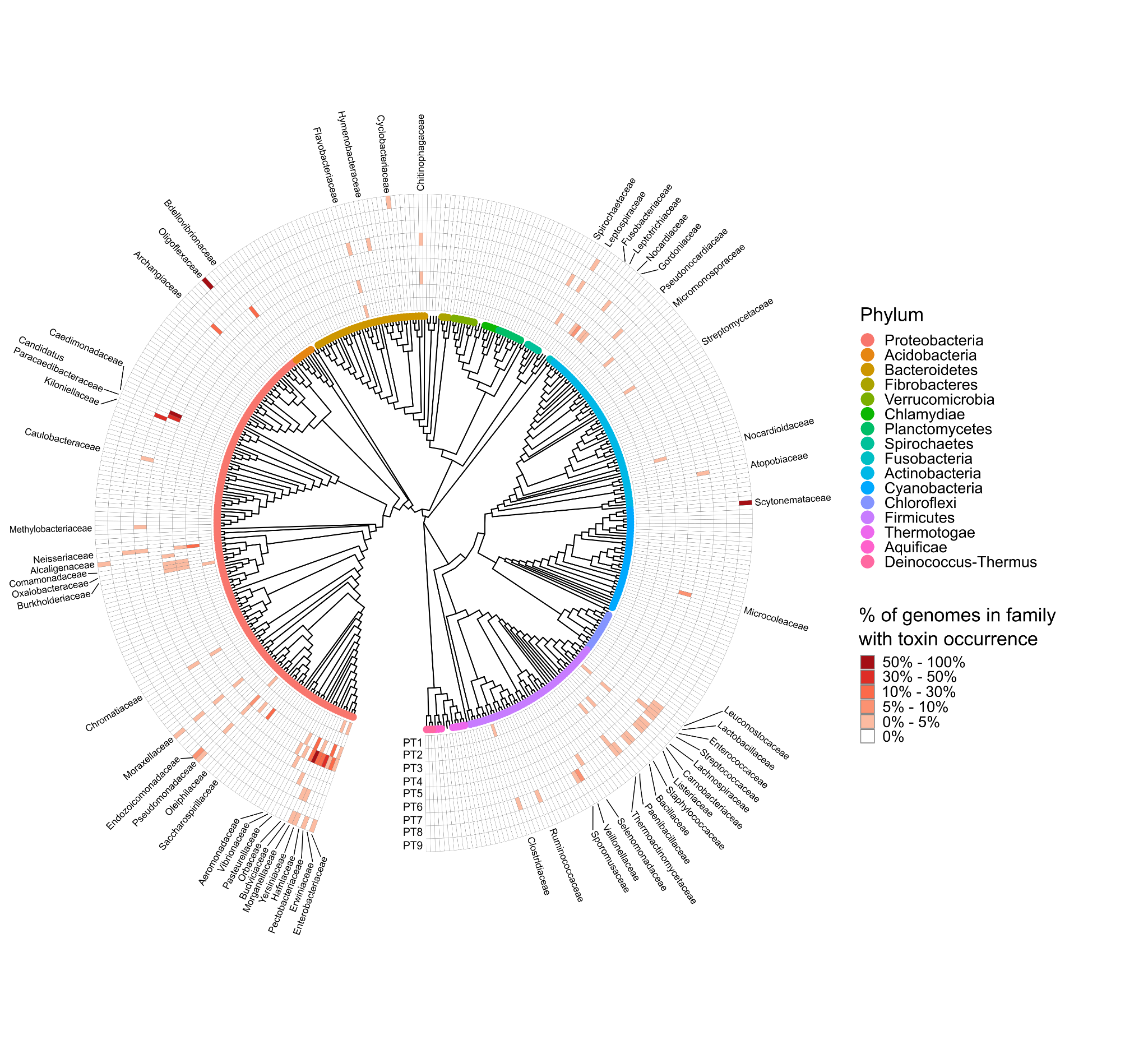

**Figure S2. Taxonomic distribution of the novel PTs.** For each family the abundance of the different PTs is presented. This is the same phylogenetic tree as in Fig 1C, but with family names.

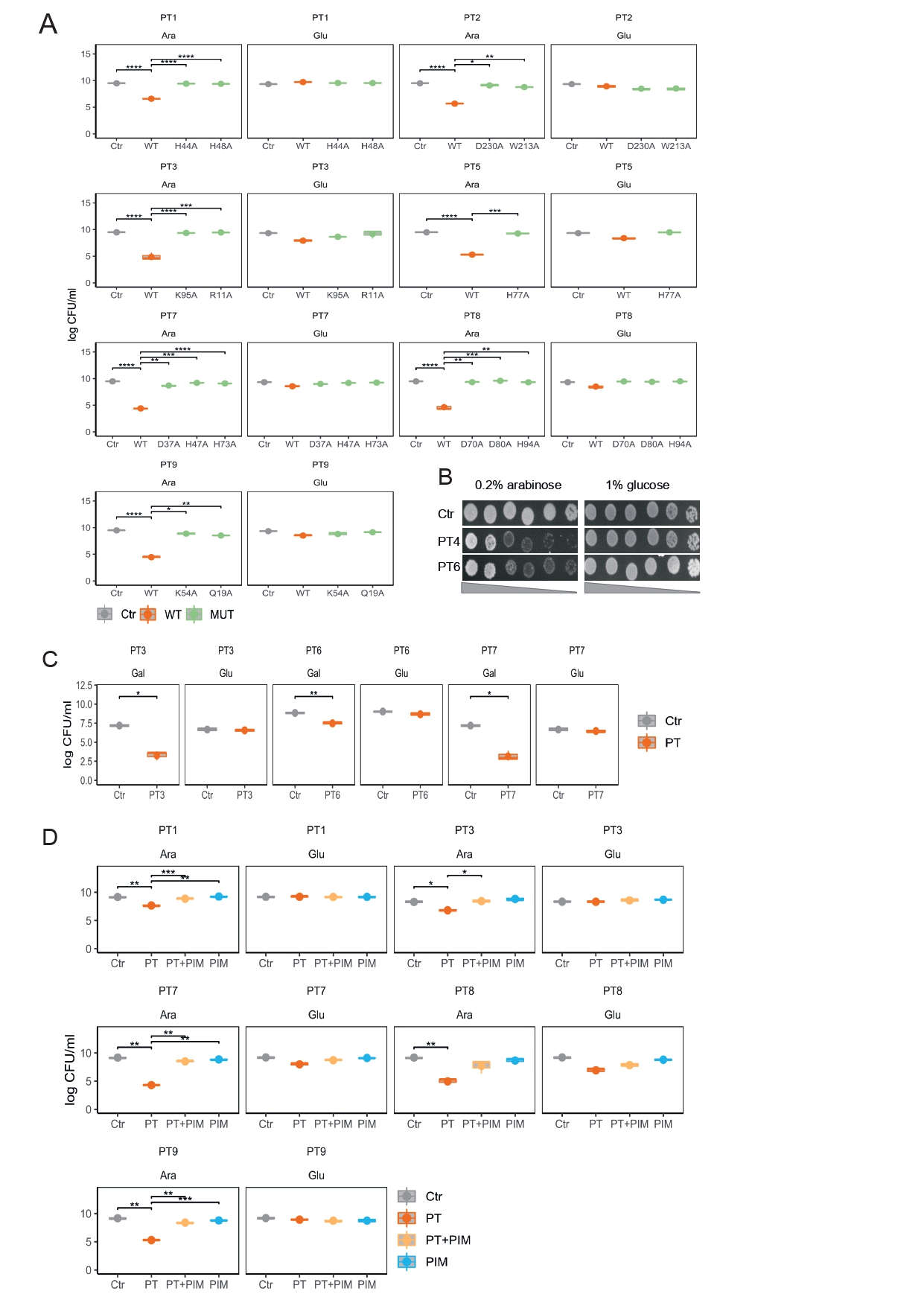

**Figure S3. The novel PTs efficiently kill bacteria and yeast. (A-B and D)** The toxins, as well as the mutants of each toxin were cloned into pBAD24 plasmieds that were transformed into *E. coli* BL21 (DE3). pBAD24 is under the control of an arabinose-inducible promoter (0.2% arabinose was used for induction). Glucose (1%) leads to repression. **(B)** Tat fused PTs for heterologous expression in *E. coli* periplasm. **(D)** The protein immunity protein (PIM) was cloned under the control of an inducible promoter system that is active only in the presence of IPTG (pET28) into *E. coli* BL21 (DE3). Boxplots represent log(CFU per mL) of **(A)** PTs and PT mutants. **(B)** Bacteria were plated in tenfold serial dilution on LB-agar in conditions that repress gene expression (1% glucose) or induce expression (0.2% arabinose). Note the bacteriostatic effect visible on spot morphology. **(D)** Toxins (PTs) and cognate immunity proteins (PIM). **(C)** PT3, PT6 and PT7 were cloned into pESC under the control of an galactose-inducible promoter *Saccharomyces cerevisiae* BY4742 and were plated in 10-fold serial dilution on SD-agar in conditions that repress gene expression (2% glucose) or induce expression (2% galactose). Boxplots represent log(CFU/mL) of yeast cells. **(A, C-D)** Asterisks indicate significant differences based on a two-sided *t-test* FDR-corrected *P* <= 0.05 (*n*=3) (* p <= 0.05, ** p <= 0.01, *** p <= 0.001). **(A-D)** As control, cells that lack the toxin (empty vector) were cloned (Ctr).

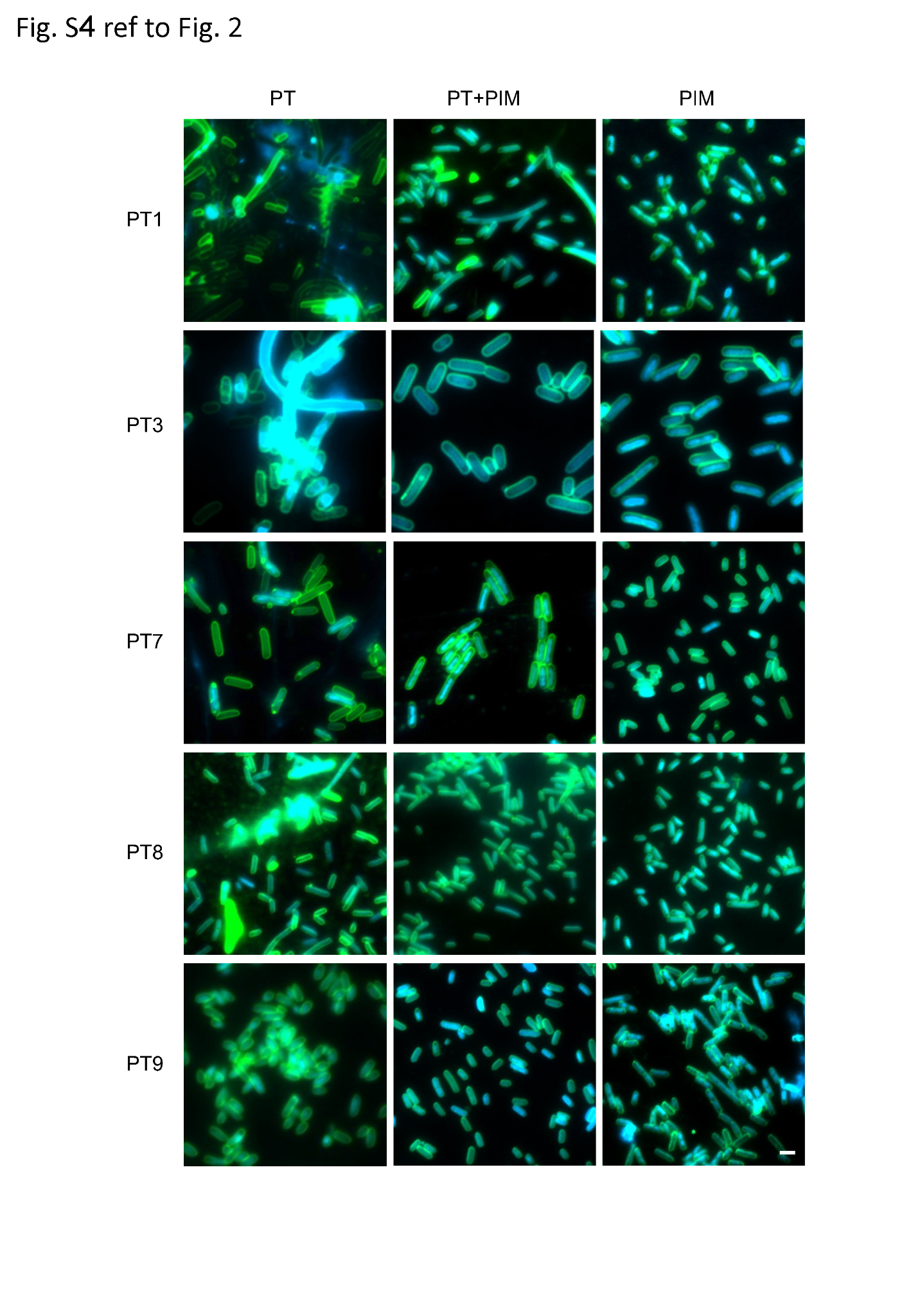

**Fig. S4. PIMs provide defense against toxicity of cognate PTs.** The toxins (PT) were cloned under the control of an arabinose-inducible promoter (pBAD24) and the immunity protein (PIM) were cloned under the control of an inducible promoter system that is active only in the presence of IPTG (pET28). Plasmids were transformed into *E. coli* BL21 (DE3). Fluorescence microscopy images of membrane stain (green) with DNA stain (blue) overlay captured at 40 min after induction with 0.2% arabinose and 0.01 mM IPTG. Each row shows cells containing one PT with its PIM. Representative images from a single replicate out of three independent replicates are shown. Scale bar corresponds to 2 μm.

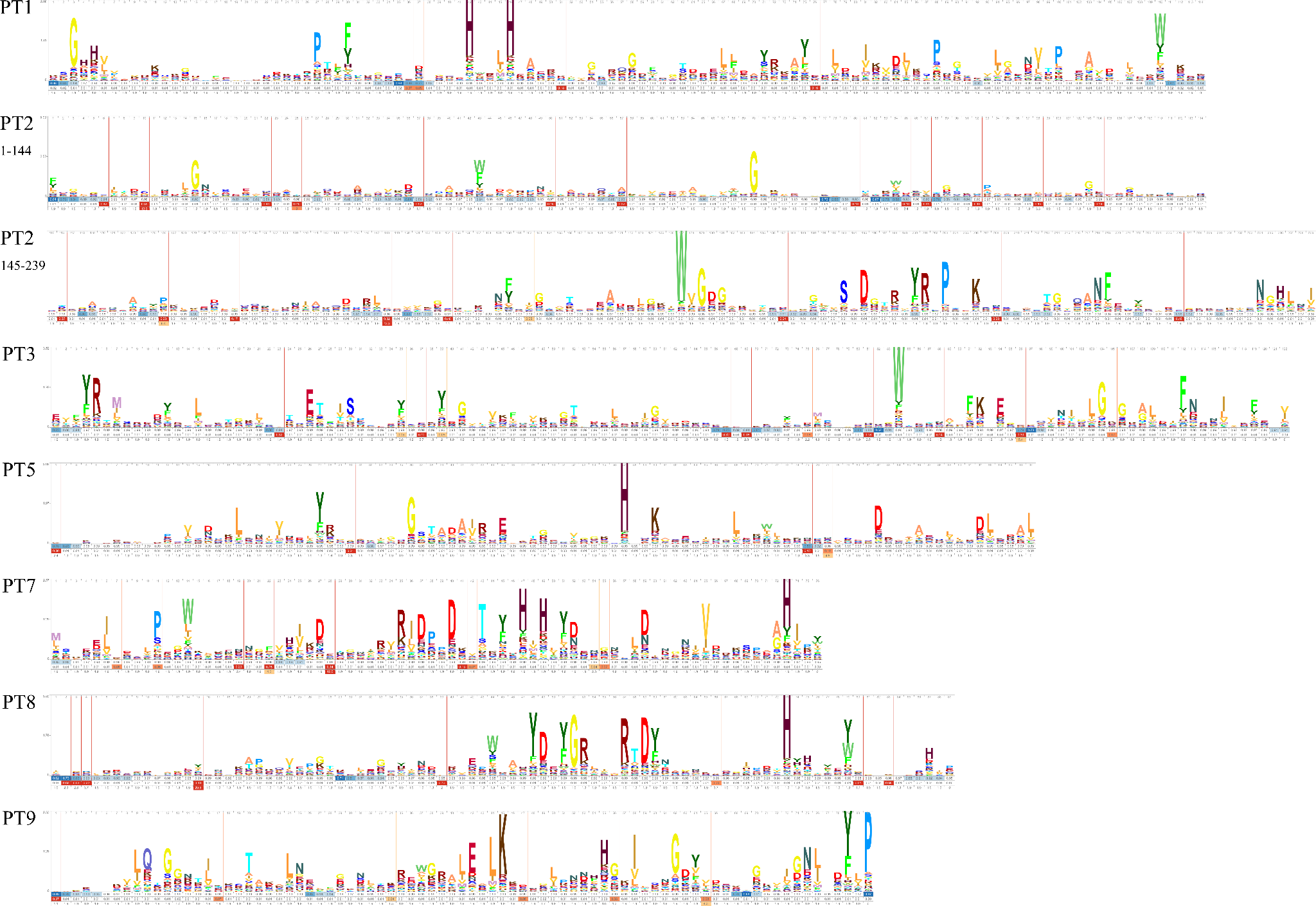

**Figure S5. Hidden Markov Models (HMM) logos for seven of the nine novel toxins.** The logos were generated out of multiple sequence alignment using skylign.org.

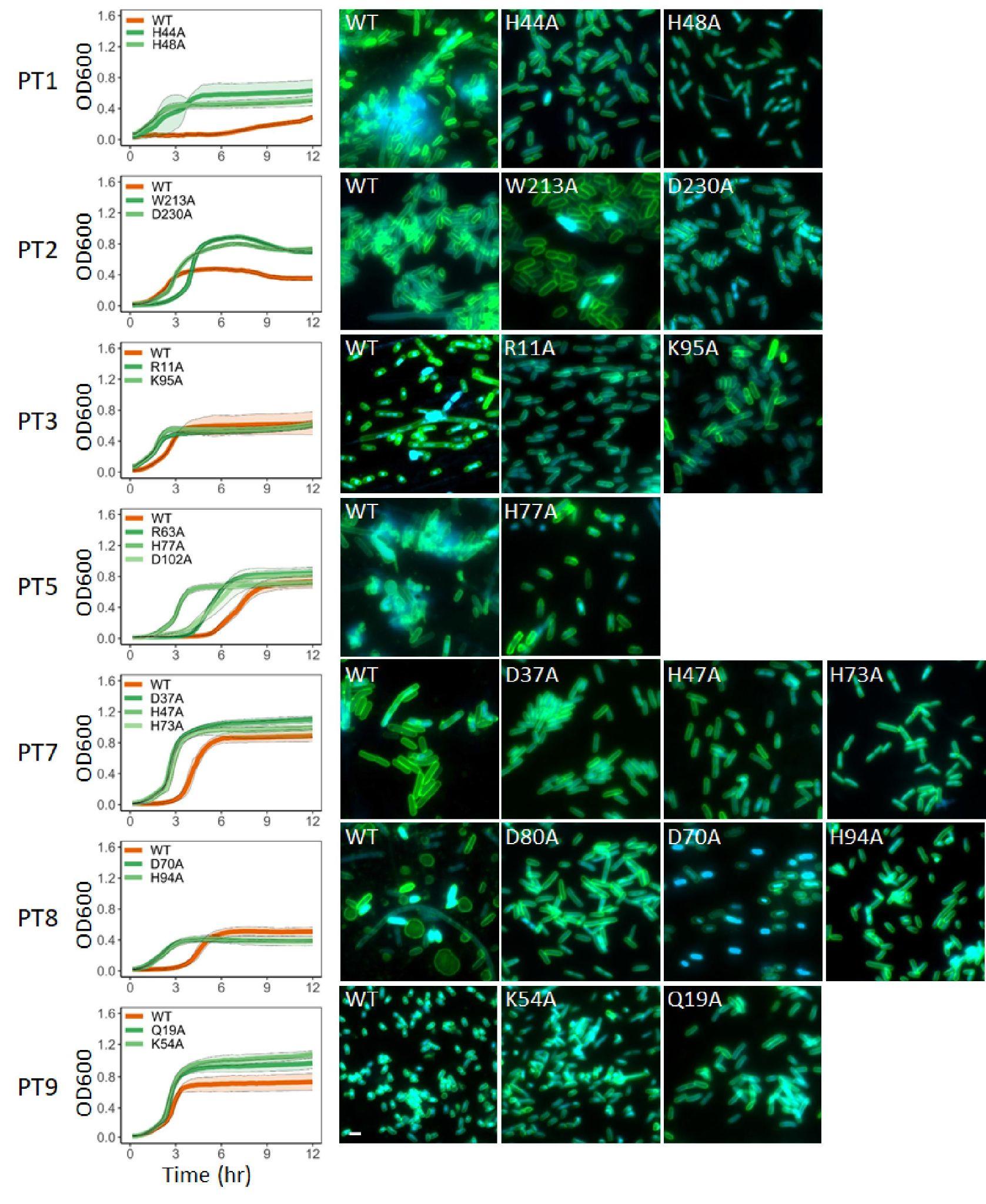

**Figure S6. Toxins lead to cell death in various manners, and mutants abrogate their cellular toxicity.** Left: The toxins, as well as the mutants of each toxin, were cloned into *E. coli* BL21 (DE3) and were grown at 37°C in LB media supplemented with 0.2% arabinose in liquid culture for 12 hours. OD600, optical density at a wavelength of 600 nm. (*n*=3 , ± SD for values). Right: for fluorescence microscopy after 40 min of induction, samples were stained with membrane and DNA stains and visualized. Shown are DNA (DAPI) (blue), membrane (FM1-43) stains overlay images of the bacterial cells with toxin or mutant toxin. Representative images from a single replicate out of three independent replicates are shown. Scale bar corresponds to 2 µm.

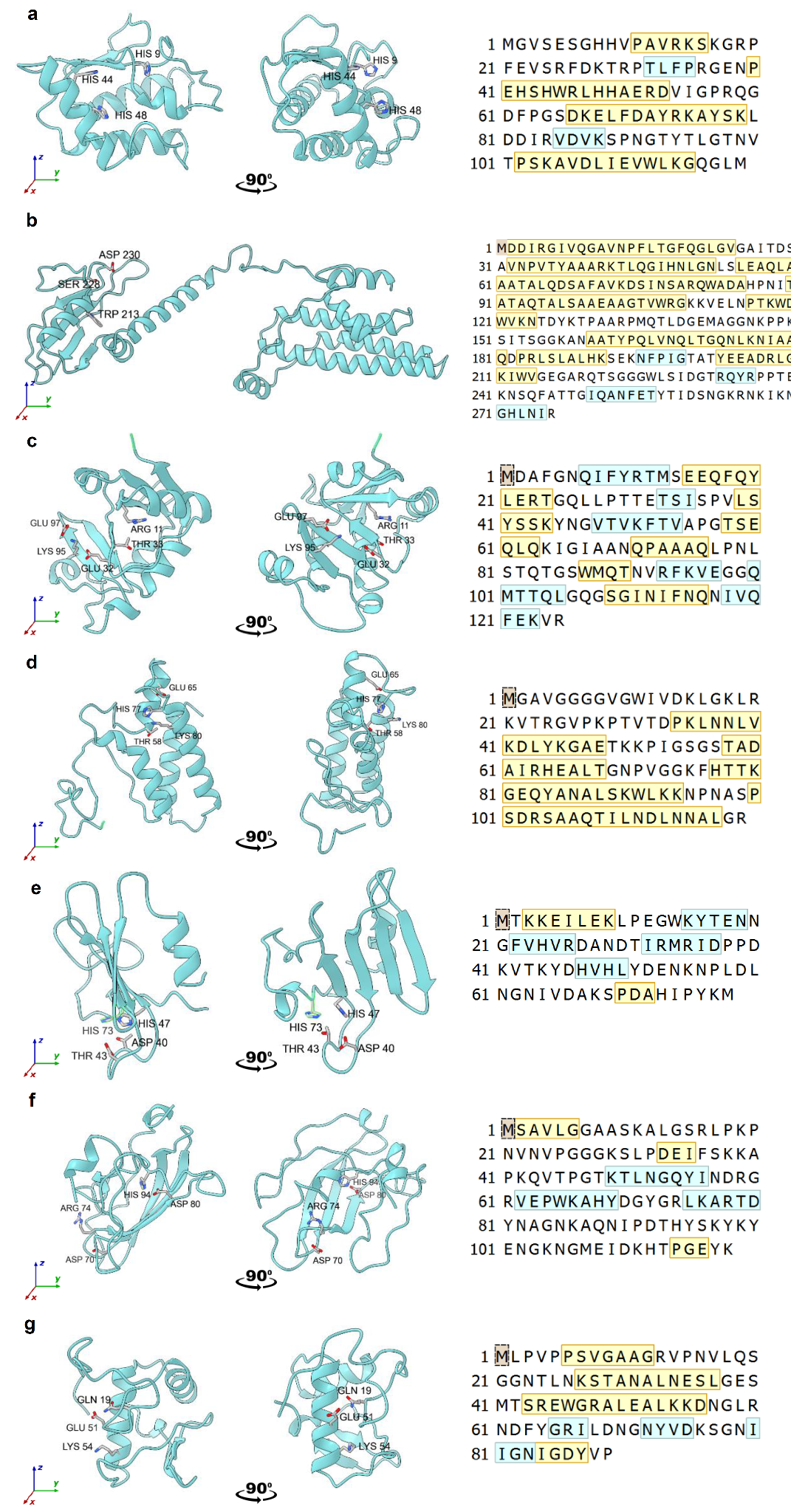

**Figure S7. PTs 3D Fold prediction**. (**A-G**) roseTTA Fold predictions for toxins PT 1-3,5,7-9 (respectively). A cartoon display using chimeraX software. From left to right: front view, side view (90° rotation to right) and protein sequence with secondary structure prediction (yellow box: 𝛼-helix, blue box: 𝛽-strand). The backbone is in cadet blue, highly conserved residues are shown as sticks. Three letter code and position number note the residue identity.

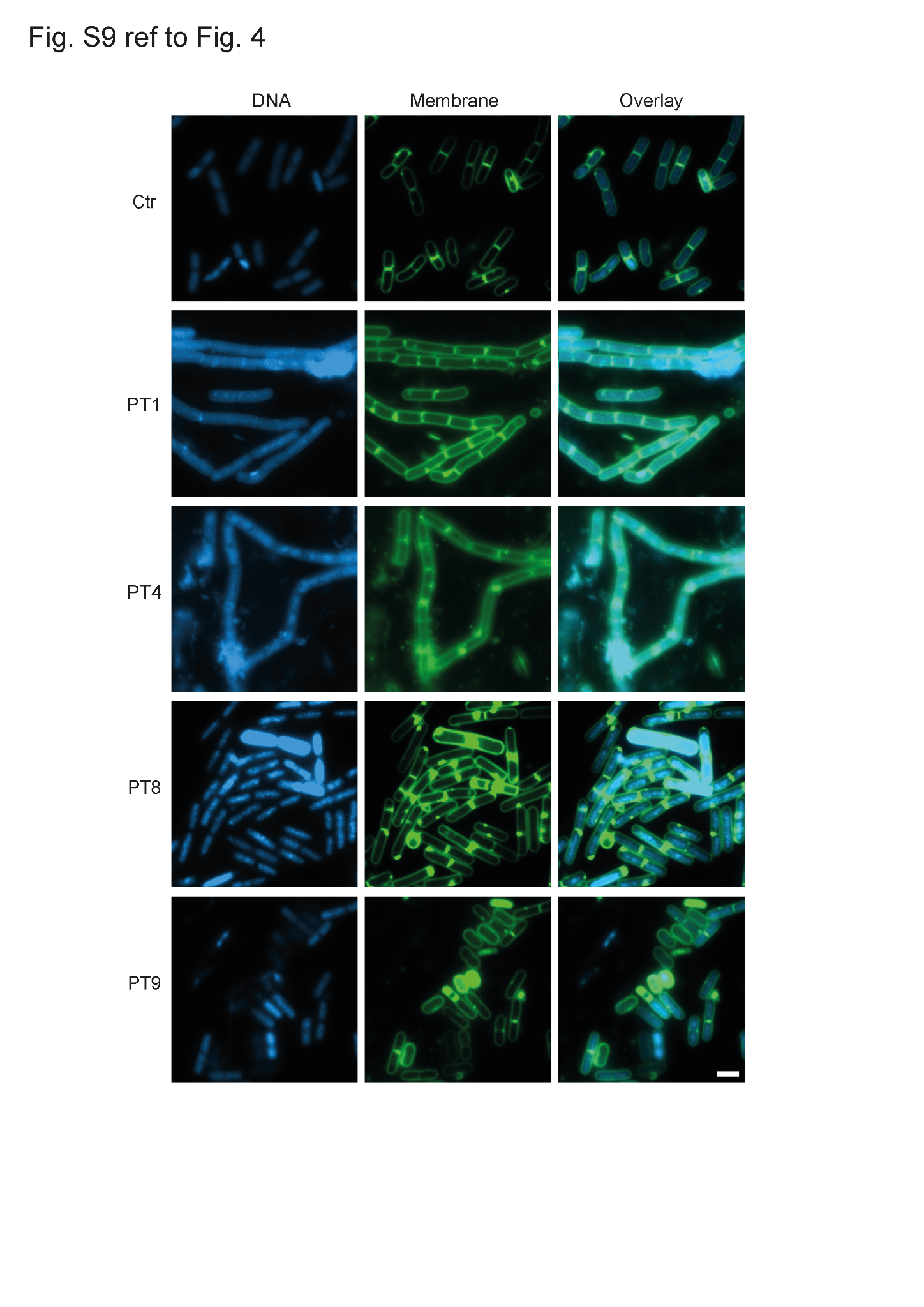

**Fig. S8. Lyasets of PT-expressing *E. coli* cells lead to *B. Subtillis* killing in various manners.**

*B. Subtills* (NCIB 3610) was grown in LB media supplement with cell lysates of PT-expressing *E. coli*. After two hours of incubation samples were stained with membrane and DNA stains and visualized by fluorescence microscopy. Shown are DNA (DAPI) (blue), membrane (FM1-43; green) and overlay images of the *B. Subtilis* cells. Ctr is the control of cell lysates of *E. coli* with an empty vector. Representative images from a single replicate out of three independent replicates are shown. Scale bar corresponds to 2 μm.

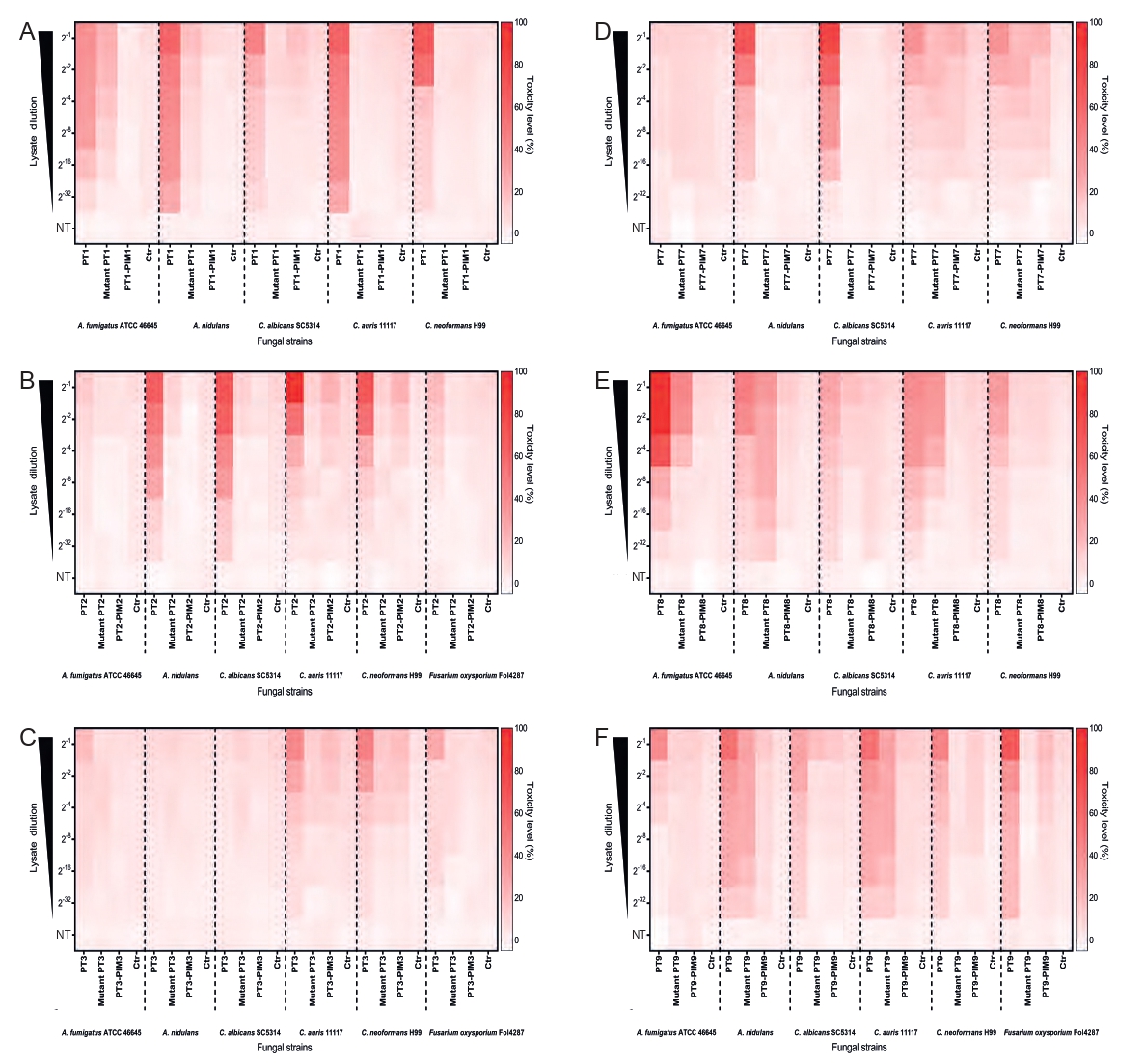

**Figure S9. Cell lysates of PT expressing *E. coli* nhibit fungal growth.**

Antifungal activity of the toxins PT1 (A), PT2 (B), PT3 (C), PT7 (D), PT8 (E), and PT9 (F) in the fungi (left to right in each panel) *A. fumigatus, A. nidulans*, *C. albicans*, *C. auris*, *C. Neoformans*, and *Fusarium oxysporum*. To determine the effect of PTs on fungal growth the indicator Alamar blue Fluorescence intensities (FI) of treated and untreated samples for each strain were obtained after 48 hours of incubation. FI were used to calculate the percentage of toxicity. The fungal strains were grown with lysates of PTs expressing *E. coli* cells. Ctr; control of lysates of *E. coli* cells with an empty vector. NT, No treatment (on y axis).

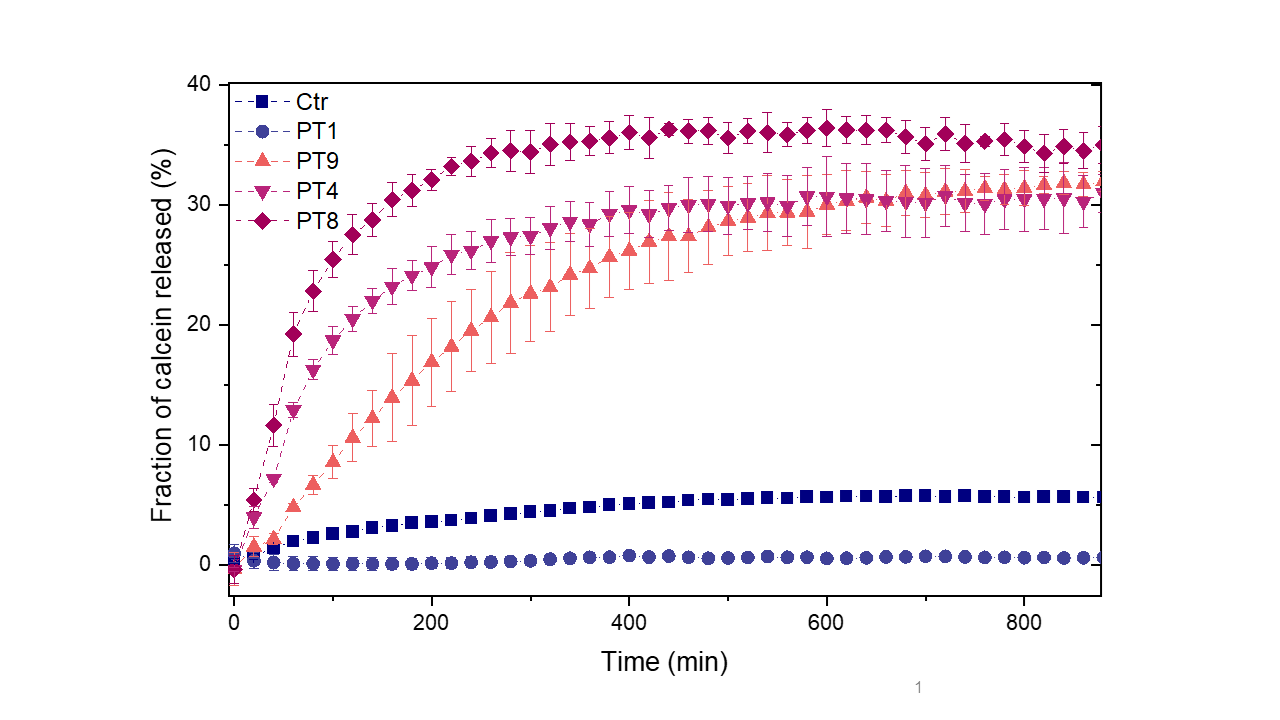
**Fig. S10. PTs permeabilize liposome membranes to elicit cell death.** Kinetic profile of calcein release from liposomes composed of *E. coli* lipid extract upon incubation with 50µl lysate of *E. coli* cells expressing PT1, PT4, PT8, PT9 and Ctr (an empty vector). Error bars represent the SEM of three technical replicates.

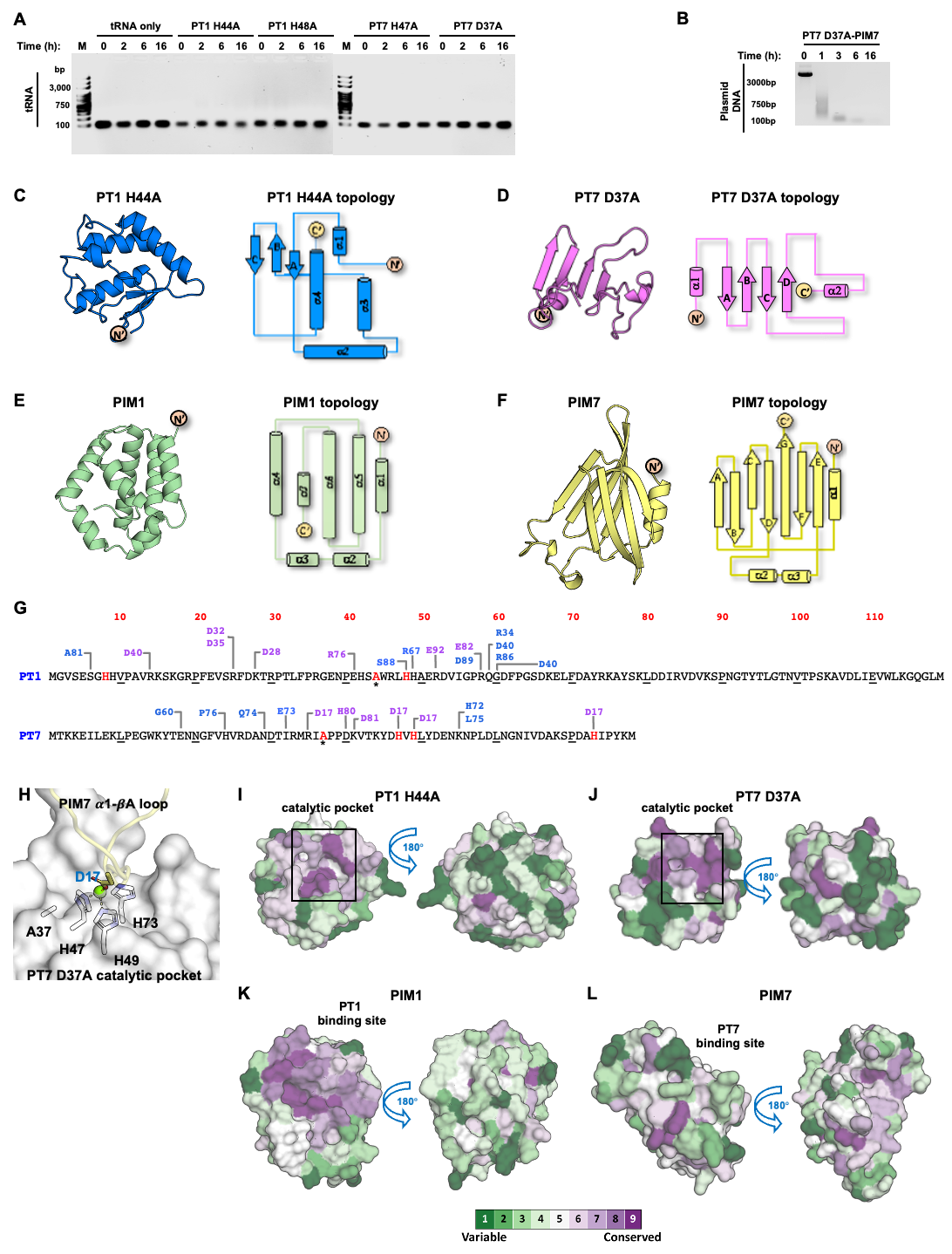

**Figure S11. Biochemical and structural characterization of PT1 and PT7 mechanism of action. (A)** In vitro nuclease activity assay of PT1 and PT7 mutants. Purified tRNA was co-incubated with buffer only (tRNA only) or with purified PT1 H44A, PT1 H48A, PT7 H47A, and PT7 D37A proteins in the presence of Mg^2+^ at 37 °C for different time points. The integrity of the tRNA was visualized on 1% agarose gel. The results indicate no ribonuclease (RNase) activity of PT1 and PT7 mutants. **(B)** Anti-toxin activity of PIM7. A purified plasmid was co-incubated with purified PT1 H48A-PIM1 complexe at 37 °C for different time points. The integrity of DNA was visualized on 1% agarose gel. No anti-toxin activity of PIM7 was observed in this assay. **(C and D)** The crystal structure of PT1 H44A **(C)** and PT7 D37A **(D)** from the PT-PIM complexes. The PTs are shown in cartoon presentation (left) and protein topology schema (right). **(E and F)** The crystal structure of PIM1 **(E)** and PIM7 **(F)** from the PT-PIM complexes. The PIMs are shown in cartoon presentation (left) and protein topology schema (right). **(G)** Schematic overview of the polar interactions between PT1 and PIM1, and PT7 and PIM7. The PT sequences are shown, and PIMs interacting residues are highlighted in purple (salt bridge) and blue (hydrogen bonds). The catalytic residues are colored in red, and an asterisk labels the mutated residues. **(H)** PT7 D37A catalytic pocket is formed by H47, H49, and H73. The interactions between PT7 D37A catalytic residues to D17, located at the tip of the PIM7 α1-βA loop, are illustrated. PT7 D37A is shown in cartoon presentation and colored in gray and PIM7 α1-βA loop is shown in cartoon presentation and colored in yellow. The side chains of the catalytic residues and PIM7 D17 are shown in sticks presentation. **(I-M)** ConSurf analysis of the PTs and PIMs. Surface presentation of PT1 H44A **(I)**, PT7 D37A **(J)**, PIM1 **(K)**, and PIM7 **(M)** with the residues colored by their conservation grades (1 to 9, where 1 is the most variable and 9 is the most conserve) indicates the high conservation of the catalytic site residues as well as the binding site of PT7.

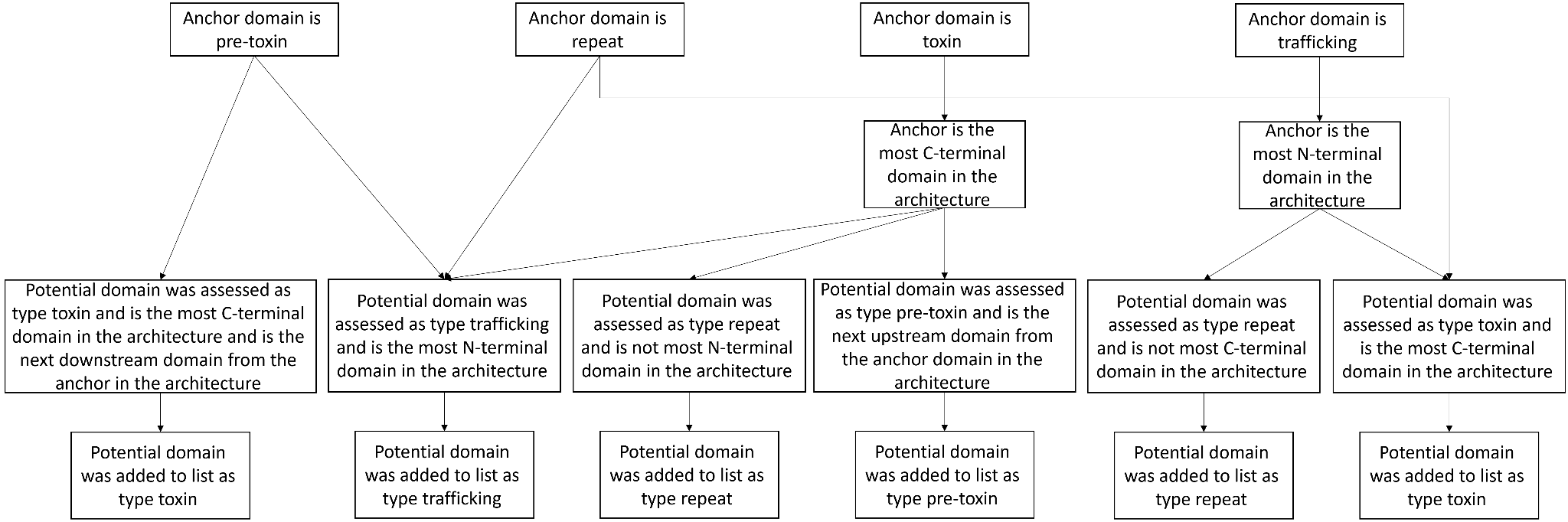

**Supplementary Figure S12**. **Schema for adding putative PT domains to the expanded list**

#### Supplementary Tables

**Table S1.** **Expanded list of 217 known and putative PT domains.** Domain types: F stands for trafficking, R for repeat, P for pre-toxin and X for toxin. Appears as a separate spreadsheet.

**Table S2. Novel PTs and PIMs identified in this study. Protein ID and existing protein annotation lay on the NCBI database.**

| **Protein Name (this work)** | **Locus Tag** | **Protein ID** | **Organism** | **Protein Length (AA)** | **Existing Protein Annotation** | **Domain Annotation** |
| --- | --- | --- | --- | --- | --- | --- |
| PT1 | C4A13_00009 | RDR26550.1 | *Escherichia marmotae* | 119 | hypothetical protein | Unannotated |
| PIM1 | C4A13_00008 | RDR26549.1 | *Escherichia marmotae* | 122 | hypothetical protein | Unannotated |
| PT2 | NMEN3081_2264 | EJU75496.1 | *Neisseria meningitidis NM3081* | 276 (the toxin should be ≤ 146 aa) | hypothetical protein | MafB at the N-terminus, the C-terminus is unannotated |
| PT3 | HTZ98_18160 | NUU72730.1 | *Ralstonia solanacearum* strain UW848 | 125 | hypothetical protein | Unannotated |
| PIM3 | HTZ98_18165 | NUU72731.1 | *Ralstonia solanacearum* strain UW848 | 157 | hypothetical protein | Unannotated |
| PT4 | EGH62_13680 | RSW81050.1 | *Klebsiella aerogenes* | 149 | adhesin | Unannotated |
| PT5 | Gene is no longer available | WP_084153669.1 | *Rheinheimera baltica DSM 14885* | 119 | hypothetical protein | Unannotated |
| PT6 | B0180_RS04825 | WP_078255899.1 | *Moraxella canis strain CCUG 8415A* | 85 | type IV secretion protein Rhs | Unannotated |
| PT7 | IE3_05452 | EJQ03417.1 | *Bacillus cereus BAG3X2-1* | 78 | hypothetical protein | Unannotated |
| PIM7 | IE3_05453 | EJQ03418.1 | *Bacillus cereus BAG3X2-1* | 117 | hypothetical protein | Unannotated |
| PT8 | LEP1GSC009_0601 | EKO85988.1 | *Leptospira interrogans serovar Grippotyphosa str. Andaman* | 118 | hypothetical protein | Unannotated |
| PIM8 | LEP1GSC009_0600 | EKO85992.1 | *Leptospira interrogans serovar Grippotyphosa str. Andaman* | 116 | hypothetical protein | Unannotated |
| PT9 | LH23_RS24110 | WP_156108067.1 | *Cedecea neteri strain M006* | 88 | hypothetical protein | Unannotated |
| PIM9 | LH23_RS23680 | WP_071842756.1 | *Cedecea neteri strain M006* | 147 | DUF4279 domain-  containing protein | DUF4279 |

**Table S3. Candidates identified by the algorithm and failed experimental validation.** Protein ID and existing protein annotation lay on the NCBI database.

| **Locus Tag** | **Protein ID** | **Organism** | **Protein Length (AA)** | **Existing Protein Annotation** | **Domain Annotation** |
| --- | --- | --- | --- | --- | --- |
| GLAD_01339 | KFC95841.1 | *Leclercia adecarboxylata* | 217 | hypothetical protein | Unannotated |
| ED901_RS00065 | WP_122094352.1 | *Rahnella sp. Larv3_ips* | 98 | hypothetical protein | Unannotated |
| I6G76_RS14020 | WP_000243841.1 | *Bacillus cereus* | 157 | hypothetical protein | Unannotated |
| LFZ28_RS26420 | WP_162802307.1 | *Salmonella enterica* | 77 | hypothetical protein | Unannotated |
| GCM10010207_38200 | GGT34515.1 | *Streptomyces atratus* | 231 | hypothetical protein | Rhs_assc_core (TIGR03696) at the N-terminus, the C-terminus is unannotated |

**Table S4. Copy number of architectures**

| **Architecture*** | **Occurrences overall** | **Occurrences with matching downstream**  **immunity protein** |
| --- | --- | --- |
| 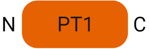 | 97 | 9 |
| 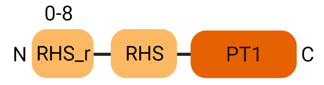 | 27 | 5 |
| 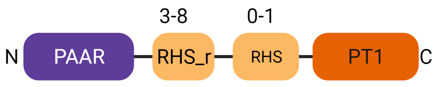 | 22 | 13 |
| 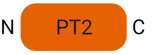 | 23 | NA |
| 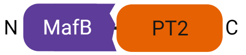 | 23 | NA |
| 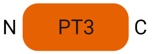 | 408 | NA |
| 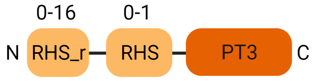 | 192 | NA |
| 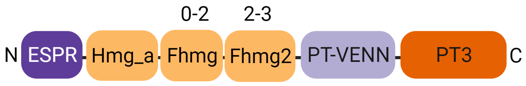 | 17 | NA |
| 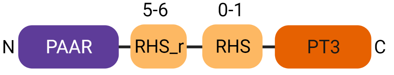 | 16 | NA |
| 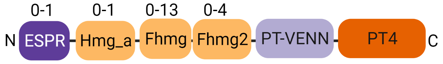 | 505 | NA |
| 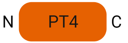 | 452 | NA |
| 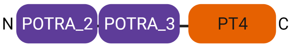 | 13 | NA |
| 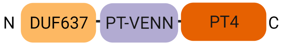 | 8 | NA |
| 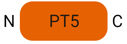 | 134 | NA |
| 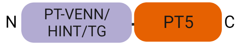 | 18 | NA |
| 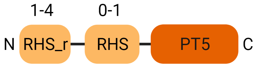 | 6 | NA |
| 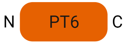 | 236 | NA |
| 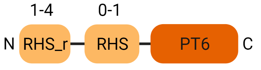 | 16 | NA |
| 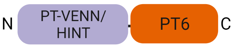 | 4 | NA |
| 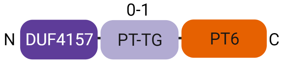 | 3 | NA |
| 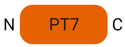 | 234 | 22 |
| 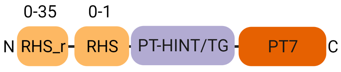 | 38 | 8 |
| 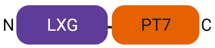 | 9 | 7 |
| 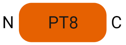 | 14 | 8 |
| 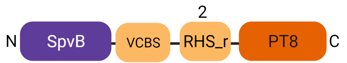 | 2 | 2 |
| 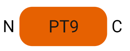 | 116 | 6 |
| 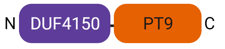 | 53 | 48 |
| 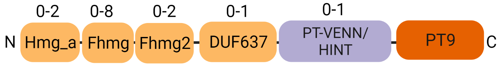 | 18 | 4 |
| 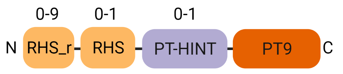 | 9 | 0 |

* The toxin domains (PTs) are shown in orange, cognate immunity genes are in blue, repeat domains are in beige, pre-toxin domains are in light purple, and trafficking domains are in dark purple. Number above the domain is the range of possible consecutive occurrences of the domain in this type of architecture.

**Table S5. All homologous proteins of PT1-9, PIM1, 3, 7-9.** Proteins were searched using DIAMOND with e value = 0.001, query cover >= 80%, “very sensitive” search mode.

**Table S6. Bacterial pathogens that encode the novel PTs.** Human pathogens are colored in red, non-human animal pathogens are in purple, plant pathogens are in green.

| **Species** | **Toxins encoded** |
| --- | --- |
| *Acinetobacter baumannii* | PT3, PT7 |
| *Actinobacillus equuli* | PT4 |
| *Aeromonas eucrenophila* | PT4 |
| *Bacillus cereus* | PT7 |
| *Bacillus mycoides* | PT7 |
| *Bacillus thuringiensis* | PT7 |
| *Brenneria alni* | PT4 |
| *Brenneria nigrifluens* | PT4 |
| *Burkholderia cenocepacia* | PT1 |
| *Burkholderia multivorans* | PT1 |
| *Clostridium botulinum* | PT1 |
| *Cronobacter sakazakii* | PT1,PT3,PT4 |
| *Dickeya zeae* | PT1,PT4 |
| *Edwardsiella anguillarum* | PT4 |
| *Edwardsiella ictaluri* | PT4 |
| *Klebsiella aerogenes* | PT3,PT4 |
| *Klebsiella pneumoniae* | PT4 |
| *Klebsiella variicola* | PT4 |
| *Leptospira interrogans* | PT8 |
| *Listeria monocytogenes* | PT7 |
| *Neisseria gonorrhoeae* | PT2 |
| *Neisseria meningitidis* | PT2 |
| *Pantoea agglomerans* | PT4 |
| *Photorhabdus luminescens* | PT4 |
| *Pluralibacter gergoviae* | PT1 |
| *Pseudomonas aeruginosa* | PT3 |
| *Ralstonia solanacearum* | PT3 |
| *Salmonella enterica* | PT3,PT4 |
| *Serratia marcescens* | PT4 |
| *Vibrio vulnificus* | PT5 |
| *Xenorhabdus cabanillasii* | PT9 |
| *Yersinia pestis* | PT4 |
| *Yersinia pseudotuberculosis* | PT4 |
| *Yersinia ruckeri* | PT4 |

**Table S7. Loci of PT3 homologs in *Pseudomonas aeruginosa* genomes from IMG.** Located as a separated spreadsheet.

**Table S8. Correlation of bacteria encoding the novel toxins with habitats and bacterial life-styles.**

| **Toxin** | **Category** | **Trait** | **Number of toxin encoding genomes with trait** | **Fisher Test Odds ratio (enrichment)** | **Fisher test q-value**  **(-log10)** | **Scoary q-value**  **(-log10)** |
| --- | --- | --- | --- | --- | --- | --- |
| PT3 | Ecosystem | Host associated | 135 | 5.085 | 15.886 | 1.814 |
|  | Ecosystem subtype | Sputum | 19 | 15.416 | 13.498 | 4.47 |
|  | Gram staining | Gram negative | 44 | 10.008 | 13.801 | 2.143 |
|  | Habitat | Human airways | 3 | 22.644 | 1.461 | 1.458 |
|  | Ecosystem type | Excretory system | 5 | 5.496 | 1.708 | 5.111 |
|  | Ecosystem type | Respiratory system | 20 | 5.763 | 6.947 | 5.476 |
| PT4 | Gram staining | Gram negative | 136 | inf | 70.182 | 3.311 |
|  | Ecosystem | Host associated | 277 | 6.537 | 38.291 | 6.77 |
|  | Ecosystem category | Human | 156 | 2.161 | 9.622 | 1.713 |
|  | Ecosystem category | Insecta | 9 | 3.666 | 2.136 | 1.55 |
|  | Ecosystem category | Fish | 7 | 3.912 | 1.84 | 1.615 |
|  | Ecosystem type | Nematoda | 3 | 8.332 | 1.391 | 2.164 |
|  | Motility | Motile | 50 | 9.455 | 9.223 | 1.716 |
| PT5 | Ecosystem | Environmental | 10 | inf | 3.66 | 1.505 |
| PT9 | Ecosystem | Host associated | 33 | 9.482 | 5.273 | 2.651 |
|  | Gram staining | Gram negative | 49 | inf | 25.219 | 1.806 |

**Table S9.** **Top DALI hits (‘Matches against PDB25’ section in DALI results) of the nine novel toxins’ predicted structures.**

| **Toxin** | **Chain (PDB)** | **Hit Protein Description** | **DALI Z-score** | **RMSD** | **Percentage of Structurally Equivalent Residues (Toxin, Hit Protein)** | **Number of Structurally Equivalent Residues** | **Length of Hit Protein (aa)** |
| --- | --- | --- | --- | --- | --- | --- | --- |
| PT1 | 7k3r-A | interferon-inducible protein Aim2 | 3.5 | 3.7 | 56.3%, 70.5% | 67 | 95 |
|  | 1ofd-A | ferredoxin-dependent glutamate synthase 2 | 3.4 | 3.7 | 64.7%, 5.2% | 77 | 1491 |
|  | 5j4a-B | trna nuclease CdiA | 3.2 | 3.1 | 56.3%, 64.4% | 67 | 104 |
| PT2 | 2uxw-A | very-long-chain specific acyl-coA dehydrogenase | 7.1 | 11.0 | 39.5%, 19.2% | 109 | 567 |
|  | 2oku-A | acyl-coa dehydrogenase family protein | 6.8 | 4.5 | 36.6%, 82.8% | 101 | 122 |
|  | 4wid-A | Rhul123 | 6.8 | 7.7 | 40.2%, 31.4% | 111 | 353 |
| PT3 | 2q5t-A | Cholix toxin | 6.3 | 3.5 | 80.8%, 16.7% | 101 | 605 |
|  | 1ddt-A | Diphtheria toxin | 6.1 | 3.3 | 77.6%, 18.5% | 97 | 523 |
|  | 6d0h-A | part: cog5654 (res domain) toxin | 5.3 | 3.8 | 76.8%, 60% | 96 | 160 |
| PT4 | 6yj4-d | NADH ubiquinone oxidoreductase chain 3 | 6.3 | 4.8 | 36.9%, 82.1% | 55 | 67 |
|  | 5xtc-V | NADH dehydrogenase [ubiquinone] iron-sulfur | 5.7 | 5.9 | 61.7%, 65.7% | 92 | 140 |
|  | 6l85-B | phosphate transporter | 5.6 | 20.5 | 50.3%, 18.7% | 75 | 401 |
| PT5 | 1oxz-A | ADP-ribosylation factor binding protein gga1 | 6.3 | 5.3 | 63.9%, 57.6% | 76 | 132 |
|  | 6swy-5 | vacuolar import and degradation protein 28 | 6.3 | 3.9 | 71.4%, 10.8% | 85 | 789 |
|  | 3jbz-A | serine/threonine-protein kinase mTOR | 6.2 | 4.8 | 68.1%, 8.4% | 81 | 960 |
| PT6 | 3psp-A | protein DOA1 | 2.8 | 3.0 | 56.5%, 15.1% | 48 | 317 |
| PT7 | 2fug-7 | NADH-quinone oxidoreductase chain 1 | 4.2 | 2.9 | 73.1%, 44.9% | 57 | 127 |
|  | 7ec1-A | glycosyl transferase, group 1 family protein | 4.1 | 2.4 | 70.5%, 10.9% | 55 | 502 |
|  | 1fj1-E | hybridoma antibody la2 (light chain) | 4.1 | 3.1 | 71.8%, 22.3% | 56 | 251 |
| PT8 | 6cp9-E | CdiA | 2.7 | 3.8 | 55.9%, 55.5% | 66 | 119 |
| PT9 | 4cht-B | DNA topoisomerase 3-alpha | 2.8 | 3.5 | 59.1%, 24.2% | 52 | 215 |
|  | 1n63-B | carbon monoxide dehydrogenase small chain | 2.8 | 3.0 | 69.3%, 7.6% | 61 | 805 |
|  | 3eps-A | isocitrate dehydrogenase kinase/phosphatase | 2.7 | 3.2 | 63.6%, 9.9% | 56 | 566 |

**Table S10. Data collection and refinement statistics for PT1 H44A – PIM1 and PT7 D37A – PIM7 complex structures.**

| Data collection | | **PT1 H44A – PIM1** | **PT7 D37A – PIM7** |
| --- | --- | --- | --- |
| Beamline | | ESRF ID-30A | ESRF ID-30A |
| Wavelength (Å) | | 0.96546 | 0.96546 |
| Space Group | | P2_1_2_1_2_1_ | P1 |
| Unit cell parameters (Å,°) | | a=48.3, b=68.0, c=69.7 | a=36.6, b=51.9, c=54.5  α=94.6, β=91.3, γ=101.6 |
| Resolution range (Å) | | 48.70-1.39 (1.41-1.39) ^a^ | 50.63-2.54 (2.58-2.54) |
| Observations | | 181,125 | 21,128 |
| Unique reflections | | 44,914 (2,282) | 12,467 (1,153) |
| Completeness (%) | | 96.4 (99.8) | 96.4 (96.8) |
| I/σ(I) | | 13.7 (2.1) | 6.6 (2.2) |
| CC_1/2_**^b^** | | 0.99 (0.76) | 0.97 (0.96) |
| R_sym_**^c^** | | 0.05 (0.60) | 0.04 (0.22) |
| R_pim_**^d^** | | 0.02 (0.32) | 0.04 (0.22) |
| Redundancy | | 4.0 (4.3) | 1.7 (1.7) |
| Refinement Statistics | | | |
| Resolution (Å) | 48.70-1.39 (1.41-1.39) | | 50.63-2.54 (2.58-2.54) |
| No. reflections | 44,847 | | 12,407 |
| R_cryst_**^e^**/R_free_**^f^** | 0.174/0.203 | | 0.214/0.260 |
| Number of atoms |  | |  |
| Protein | 3,225 | | 3,174 |
| Water | 205 | | 0 |
| Ligand (Mg^2+^) |  | | 1 |
| Average B value (Å^2^) | | | |
| All proteins | 20 | | 66 |
| Water | 31 | |  |
| Ligand (Mg^2+^) |  | | 86 |
| R.m.s. deviation from ideal geometry | | | |
| Bond length (Å) | 0.007 | | 0.012 |
| Bond angle (º) | 0.8 | | 1.4 |
| Ramachandran Plot (%) **^g^** | | | |
| Favored | 99.7 | | 97.5 |
| Outliers | 0.0 | | 0.5 |
| PDB ID | XXXX | | YYYY |

### **Table S11. PCR primers used for site-directed mutagenesis and for PTs-*gfp* fusion**

| **Product** | **purpose** | **primers** | **Abolishing**  **toxicity*** |
| --- | --- | --- | --- |
| pBAD_PT1_H44A | Inverse PCR | PT1_H44A_F GGAGCACTCAGCGTGGCGTCTGCACCACG  PT1_H44A_R GGGTTTTCACCACGGGGA | +++ |
| pBAD_PT1_H48A | Inverse PCR | PT1_H48A_F GCGCACGCAGAGCGCGATG  PT1_H48A_R CAGACGCCAATGTGAGTGC | +++ |
| pBAD_PT2_W213A | Inverse PCR | PT2_W213A_F GGGAAAAATTGCGGTCGGAGAAGGTGC  PT2_W213A_R AGACGGTCCGCTTCCTCA | +++ |
| pBAD_PT2_D230A | Inverse PCR | PT2_D230A_F GCTGAGCATCGCGGGCACCCGCC  PT2_D230A_R CAGCCGCCACCCGAAGTT | +++ |
| pBAD_PT3_R11A | Inverse PCR | PT3_R11A_F AATTTTCTATGCGACTATGTCGGAAGAACAGTTCCAATACCTG  PT3_R11A_R TGGTTACCGAACGCGTCC | +++ |
| pBAD_PT3_K95A | Inverse PCR | PT3_K95A_F TGTTCGTTTCGCGGTGGAAGGCGGTC  PT3_K95A_R TTGGTCTGCATCCAGCTC | +++ |
| pBAD_PT5_R63A | Inverse PCR | PT5_R63A_F TGATGCAATTGCGCATGAGGCGCTCACCG  PT5_R63A_R GCTGTCGAACCACTCCCG | - |
| pBAD_PT5_H77A | Inverse PCR | PT5_H77A_F TGGCAAATTTGCGACGACGAAAGGGGAACAGTATGC  PT5_H77A_R CCGACTGGATTGCCGGTG | +++ |
| pBAD_PT5_D102A | Inverse PCR | PT5_D102A_F GTCTCCGTCCGCGCGCAGCGCAG  PT5_D102A_R GCATTAGGATTTTTCTTAAGCCACTTACTC | - |
| pBAD_PT7_D37A | Overlapping fragments | PT7_NCO_R GAGGATCCCCGGGTACCATGGGTCGATCAGCGTACG  PT7_NCO_F TGGGCTAGCAGGAGGAATTCATGACTAAGAAGGAGATTCTGG  PT7_D37_NEB_F ATGCGTATTGCGCCGCCAGACAAGGTTAC  PT7_D37A_NEB_R TGTCTGGCGGCGCAATACGCATGCGAATCGTG | +++ |
| pBAD_PT7_T43A | Overlapping fragments | PT7_T43A_F AGACAAGGTTGCGAAGTACGACCACGTTC  PT7_T43A_R TGGTCGTACTTCGCAACCTTGTCTGGCGGATC | + |
| pBAD_PT7_H47A | Overlapping fragments | PT7_H47A_F AAGTACGACGCGGTTCACCTGTACGATGAAAAC  PT7_H47A_R CAGGTGAACCGCGTCGTACTTCGTAACCTTGTCTG | +++ |
| pBAD_PT7_H73A | Overlapping fragments | PT7_H73A_NEB_F CCCGGACGCCGCGATCCCGTACAAGATGTGATGAC  PT7_H73A_NEB_R ACGGGATCGCGGCGTCCGGGCTCTTG | +++ |
| pBAD_PT8_D70A | Overlapping fragments | PT8_PBAD_R AGGATCCCCGGGTACCATGGTCACTTGTACTCACCCGG  PT8_PBAD_F TGGGCTAGCAGGAGGAATTCATGTCTGCAGTACTGGGC  PT8_H70A_R GTCCGTACCCCGCGTAGTGGGCCTTCCACG  PT8_H70A_F GGCCCACTACGCGGGGTACGGACGCTTGAAG | +++ |
| pBAD_PT8_D80A | Inverse PCR | PT8_D80A_F GGCTCGCACCGCGTACAACGCGG  PT8_D80A_R TTCAAGCGTCCGTACCCA | +++ |
| pBAD_PT8_H94A | Overlapping fragments | PT8_H94A_NEB_R ACTTAGAATACGCGGTGTCTGGAATATTCTGCGC  PT8_H94A_F CCAGACACCGCGTATTCTAAGTATAAGTACGAAAACGGC | +++ |
| pBAD_PT9_Q19A | Overlapping fragments | PT9_PBAD_F TTGGGCTAGCAGGAGGAATTCATGTTGCCAGTGCCACC  PT9_PBAD_R CTCTAGAGGATCCCCGGGTACCATGGTTATGGCACGTAGTCCCCG  PT9_Q19A_F GAACGTGTTGGCGTCTGGCGGCAACACTC  PT9_Q19A_R CCGCCAGACGCCAACACGTTCGGCACACG | +++ |
| pBAD_PT9_T29A | Overlapping fragments | PT9_T29A_R GCATTCGCCGCGCTTTTATTCAGAGTGTTGCCG  PT9_T29A_F CTGAATAAAAGCGCGGCGAATGCGCTGAACG | + |
| pBAD_PT9_K54A | Inverse PCR | PT9_K54_F GAGGCCCTGGCGAAGGACAACGGTCTCCG  PT9_K54_R TGTCCTTCGCCAGGGCCTCAAGCGC | +++ |
| pBAD_sfGFP6XhIS_F |  | **sfGFP6XhIS_F**  ATGCGTAAAGGCGAAGAG  **sfGFP6XhIS_R**  TCTAGAGGATCCCCGGGTACTTAGTGATGGTGATGGTG | —- |
| pBAD_PT1*-sf-gfp-his* |  | **PT1-F**  GGCTAGCAGGAGGAATTCACATGGGGGTTTCTGAATCGG  **PT1-gfp-R**  **AGCTCTTCGCCTTTACGCATCATCAGGCCCTGACCCTTC** | +++ |
| pBAD_PT7-*sf-gfp-his* |  | **PT7-F**  GGCTAGCAGGAGGAATTCACATGACTAAGAAGGAGATTCTG  **PT7-gfp-R**  AGCTCTTCGCCTTTACGCATCATCTTGTACGGGATGTG | +++ |
| pBAD_PT8-*sf-gfp-his* |  | **PT8-F**  GGCTAGCAGGAGGAATTCACATGTCTGCAGTACTGGGC  **PT8-gfp-R**  **AGCTCTTCGCCTTTACGCATCTTGTACTCACCCGGGGTG** | +++ |
| pBAD_PT9-*sf-gfp-his* |  | **PT9-F**  **GGCTAGCAGGAGGAATTCACATGTTGCCAGTGCCACCG**  **PT9-gfp-R**  **AGCTCTTCGCCTTTACGCATTGGCACGTAGTCCCCGATG** | +++ |

*+ minor effect; +++ complete abolishment of activity; - no effect

**Table S12 Bacterial and fungal strains**

| **Strain** | **genotype** | **use** |
| --- | --- | --- |
| *E.coli* sp. DH5𝛼 | F^–^ *endA1* *glnV44* *thi-1* *recA1* *relA1* *gyrA96* *deoR* *nupG* *purB20* φ80d*lacZ*ΔM15 Δ(*lacZYA-argF*)U169, hsdR17(*r_K_*^–^*m_K_*^+^), λ^–^ | Plasmid propagation |
| *E. coli* sp. BL21 (DE3) | B F^–^ *ompT* *gal* *dcm* *lon* *hsdS_B_*(*r_B_*^–^*m_B_*^–^) λ(DE3 [*lacI* *lacUV5*-*T7p07* *ind1* *sam7* *nin5*]) [*malB*^+^]_K-12_(λ^S^) | Het. Expression |
| Baking yeast sp. BY4742 | MATα His3Δ1 leu2Δ0 lys2Δ0 ura3Δ0 | Het. Expression |
| *B. Subtills* (NCIB 3610) | Wild type | Antibacterial experiments |
| *E. coli* MG1655 imp4213 | Wild type | Antibacterial experiments |
| *Aspergillus fumigatus ATCC 46645* | Wild type | Antifungal experiments |
| *Aspergillus nidulans* | Wild type | Antifungal experiments |
| *Candida auris 11117* | Wild type | Antifungal experiments |
| *Candida albicans SC5314* | Wild type | Antifungal experiments |
| *Criptococcus neoformans H99* | Wild type | Antifungal experiments |
| *Fusarium oxysporium Fol4287* | Wild type | Antifungal experiments |
